# RNA and FUS act in concert to prevent TDP-43 spatial segregation

**DOI:** 10.1101/2023.07.25.550456

**Authors:** Clément Demongin, Samuel Tranier, Vandana Joshi, Léa Ceschi, Bénédicte Desforges, David Pastré, Loïc Hamon

## Abstract

FUS and TDP-43 are two self-adhesive aggregation-prone mRNA-binding proteins whose pathological mutations have been linked to neurodegeneration. While TDP-43 and FUS form reversible mRNA-rich compartments in the nucleus, pathological mutations promote their respective cytoplasmic aggregation in neurons with no apparent link between the two proteins except their intertwined function in mRNA processing. By combining analyzes in cellular context and at high-resolution *in vitro*, we unraveled that TDP-43 is specifically recruited in FUS assemblies to form TDP-43 rich sub-compartments but without reciprocity. The presence of mRNA provides an additional scaffold to promote the mixing between TDP-43 and FUS. Accordingly, we also found that the pathological truncated form of TDP-43, TDP-25, which has an impaired RNA binding ability, no longer mixes with FUS. Together, these results suggest that the binding of FUS along nascent mRNAs enables TDP-43, which is highly aggregation-prone, to mix with FUS phase in order to form mRNA-rich sub-compartments. A functional link between FUS and TDP-43 may explain their common implication in Amyotrophic Lateral Sclerosis (ALS).

## Introduction

TDP-43 and FUS have been under scrutiny over the last past years due to their link with amyotrophic lateral sclerosis (ALS) and frontotemporal lobar degeneration (FTLD). Both FUS and TDP-43 are nuclear proteins that assemble into insoluble cytoplasmic aggregates in the neurons of patients affected by these two incurable neurodegenerative diseases.[1–4]. They also share similar targets and structure, both proteins being associated to mRNA and harboring long unstructured self-adhesive domains to regulate their higher order assemblies. A tight regulation of dynamic assemblies of TDP-43 and FUS is required to fulfill their intertwined functions associated to the mRNA life cycle: transcriptional regulation, pre-mRNA splicing, mRNA localization and processing [5–7]. To enable reversibility of theses assemblies, TDP-43 and FUS should remain soluble which is a challenging task for cells due to the presence of self-adhesive Low Complexity Domains (LCD) prone to aggregation. The solubility of TDP-43 and FUS is also controlled by their binding to RNA, which is their main partner *in vivo*. Indeed, a high RNA/RBP molar ratio favors the dynamics and reversible assemblies of TDP-43 and FUS in the nucleus under physiological conditions [8, 9]. A high RNA/RBP molar ratio could be also observed in the cytoplasm in mRNA-rich stress granules after acute cellular stress [10]. The presence of TDP-43 and FUS in stress granules may indeed preserve their solubility [9, 11]. Conversely, a low RNA/RBP ratio promotes multivalent protein-protein interactions, mainly via LCDs, and leads to the formation of insoluble aggregates. In an alternative model, stress granules can be considered as crucibles in the formation of cytoplasmic inclusions linked to neurodegenerative diseases [12, 13] due to the high concentration of LCD-rich proteins that promotes the formation of an aggregation-prone sub-compartment. Again in agreement with a higher protein solubility in the presence of mRNA, an altered binding between these proteins and RNA, in particular through pathological mutations, can lead to their aggregation [14, 15].

Their intertwined functions in the regulation of transcription and in mRNA splicing lead TDP-43 and FUS to associate with approximately 30% of the transcriptome in the human brain [16]. However, their binding sites on mRNAs, as observed by cross-linking and immunoprecipitation (CLIP) methods, seem quite distinct [17, 18]. FUS binds nonspecifically along nascent mRNAs [19] while TDP-43 rather forms clusters in GU-rich sequences. Despites these differences, the sets of genes with altered expression levels upon TDP-43 or FUS knockdown exhibit significantly overlapping transcriptome profile [20]. In addition, their coordination is also necessary to regulate the expression of common targets such as histone deacetylase 6 (HDAC6) [21]. In animal models like drosophila and zebrafish, combination of knockdown, overexpression, and rescue studies of TDP-43 and FUS supports the notion that TDP-43 and FUS participate to mRNA maturation pathways [22–24]. Beyond mRNA-related functions, FUS and TDP-43 have also common protein partners. For example, both of them have the ability to stall RNA PolII processing [25, 26]. Finally, analyzes by co-immunoprecipitation demonstrate that these two proteins could be present in their mutual interactome [27–29] while they almost never coexist in cytoplasmic aggregates in neurons of ALS/FTLD patients [30, 31]. There are thus numerous lines of evidence indicating the simultaneous presence of FUS and TDP-43 on the same RNA substrates at the same time to perform similar or complementary biological functions.

We propose to study the original relationship between these two RBPs, particularly their interplay for the binding to mRNA and the consequence of these interactions in their ability to form compartments in which they could mix or demix. Our working hypothesis is based on the low specificity of the FUS for RNA sequences [19, 32]. FUS rather show a preference for RNA structures [33, 34] with an ability to remodel RNA via its interaction with RGG domains [35]. Through these characteristics, FUS may bind to nascent mRNAs to form FUS-rich compartments in which TDP-43 is recruited and may bind to intron-rich GU sequences with high affinity via its RNA recognition motifs (RRM) [17, 36, 37]. The binding of TDP-43 to RNA is cooperative, which, along with self-attracting N-terminal domain, can secure the formation of TDP-43-rich compartments on GU-rich sequences on long RNAs [38]. Here, we combine analyzes in a cellular context using microtubules as nano-platforms [39] with structural information obtained at the single molecule scale [40] to gain access to the spatial organization of TDP-43 and FUS. We first demonstrate that TDP-43 and FUS interact with each other. However, FUS is able to accept TDP-43 in its compartment while the opposite is not observed. *In vitro*, we evidence that the partial miscibility between FUS and TDP-43 is enhanced by the presence of RNA. We observe a homogenous distribution of TDP-43 along RNA, when TDP-43 is present at low concentration while at high concentration, TDP-43 forms sub-compartments still associated to FUS. We also demonstrate that the colocalization of FUS and TDP-43 in stress granules is more pronounced than with other RBPs but when TDP-43 loose its ability to bind cooperatively to RNA, the colocalization is affected. Finally, we observe that the RNA binding capacity of TDP-43 is essential to preserve the miscibility of TDP-43. Consistently, pathological truncations of TDP-43 having lost all or part of the RRMs are excluded from the FUS-RNA complexes and aggregate independently of FUS. Therefore, the interaction between these two ALS-linked proteins can significantly contribute to their essential functions in RNA metabolism. Importantly, the interaction TDP-43 / FUS may help to preserve TDP-43 from forming distinct assemblies on its own, which would constitute an early step toward TDP-43 aggregation.

## Materials and methods

### Cell experiments

#### Plasmid preparation for MT Bench bait/prey method

For the microtubule bench method, plasmids encoding proteins of interest fused to RFP and Microtubule-Binding Domain of Tau (MBD) were produced as previously described [39, 41] and are summarized in supplementary table 1. cDNA of full length FUS, TDP-43, were amplified using primers containing Pac1 and Asc1 restriction sites. Fragments obtained were inserted into the backbone entry plasmid RFP-MBD-pCR8/GW/TOPO previously digested with the corresponding restriction enzymes. Then recombination step was done (Gateway® LR Clonase® II Enzyme mix, INVITROGEN, cat n°11791020) in the final mammalian cellular expression plasmid pEF-Dest51 (Invitrogen^TM^). GFP-fused constructs were obtained as described previously [41]. cDNAs encoding for sequences of proteins of interest were amplified and inserted into pEGFP-N1 vector (Clontech). Each DNA construct was checked by conventional Sanger sequencing of purified plasmid.

#### Cell culture, transfection and fixation for MT Bench experiments

U2OS cells were used for the MT bench given their well-suited morphology to visualize microtubules. They were grown in DMEM (Dulbecco’s Modified Eagle’s Medium, high glucose, Sigma) with 10 % FBS and penicillin/streptomycin 100 μg/mL (GIBCO Life Technologies). Around the confluence, cells were transferred in 4-well plates on 12 mm coverslips for transfection. U2OS cells were transfected with 500 ng (1:1 ratio) of plasmids RBP1-RFP-MBD/RBP2-GFP (FUS-RFP-MBD and TDP-RFP-MBD with HuR-GFP, G3BP-GFP, YB1-GFP, Lin28-GFP, LARP6-GFP, SAM68-GFP, and U2AF-GFP) with lipofectamine 2000^TM^ reagent (Invitrogen) added to the culture medium. Medium was changed after 4 h incubation to remove lipofectamine, and then cells were incubated overnight at 37 °C under a controlled atmosphere (5 % CO2). Cells were washed with PBS and fixed, firstly with methanol 100 % at -20 °C for 20 min, then with PFA 4 % in PBS for 30 min at 37°C. Finally, cells were mounted on glass slides with MOWIOL (Sigma).

#### Observation and measurement of the correlation coefficient in the bait/prey method

Images of transfected cells were acquired with an oil immersed objective 63x/1.4 on an inverted microscope (Axiovert 220; Carl Zeiss 5 MicroImaging, Inc, Hamamatsu C10600, Axio Vision software). Exposure times were adjusted to obtain comparable fluorescence intensity between the different channels. Images were cropped for each cell and intensity and contrast parameters were adjusted with imageJ software. Correlation between RFP and GFP signals were determined as previously described [42] and process is summarized in the supplementary file S1. Briefly, images were treated with ImageJ software and intensity parameters were adjusted to highlight microtubules. A line crossing several microtubules was drawn (10 to 20 μm length) tangent and close to the nucleus and fluorescence intensities along this line were measured for both colors. A plot was generated with peaks corresponding to microtubules. If the proteins of interest interact on microtubules, overlapping of peaks for green and red signals is observed. The line profiles for each channel were transformed into numerical values. A Pearson correlation coefficient was calculated using Microscoft Excel CORREL function based on these values. For each condition 3 lines per cell and more than 10 cells were analyzed.

#### Plasmid preparation for MT Bench compartmentalization method

HuR-RFP-MBD, Sam68-RFP-MBD, FUS-GFP-MBD and TDP-43-GFP-MBD plasmids (supplementary table 2) were obtained as described previously for bait/prey method [39]. cDNA for truncated forms of TDP-43 and FUS (supplementary table 2), were amplified using designed primers containing Pac1 and Asc1 restriction sites. Fragments obtained were inserted into the backbone entry plasmid RFP-MBD-pCR8/GW/TOPO or GFP-MBD-pCR8/GW/TOPO previously digested with the corresponding restriction enzymes. Then recombination step was done (Gateway® LR Clonase® II Enzyme mix, INVITROGEN, cat n°11791020) in the final mammalian cellular expression plasmid pEF-Dest51 (Invitrogen^TM^). DNA sequencing was done for each production.

#### Cell transfection and preparation for MT Bench compartmentalization method

U2OS cells were prepared, transfected, and fixed as described previously. To study the compartmentalization of the proteins according to their concentration, cells were transfected with several amounts of plasmids of FUS-RFP-MBD/TDP-43-GFP-MBD, Sam68-RFP-MBD/FUS-GFP-MBD, Sam68-RFP-MBD/TDP-43-GFP-MBD, HuR-RFP-MBD/TDP-43-GFP-MBD, or HuR-RFP-MBD/FUS-GFP-MBD (corresponding to 0.5/5, 0.5/2.5, 0.5/1.25, 1/1, 2.5/1, 5/1, 10/1 µg/µg of plasmids). To avoid bias linked to difference in fluorescence intensities in the analysis, acquisition parameters (exposure time and exposition intensity) were adjusted to obtain comparable levels for green and red intensities. RBP1/RBP2 expression level ratios for each plasmid ratio, were recalculated using adjusted parameters: exposure time [Expo] and measured fluorescence intensity [I] for each cell were divided by the mean values of exposure time and fluorescence intensities respectively for at least 20 cells. [plasmid] corresponds to the plasmid ratio (in µg/µg) used for the transfection and is ranging from 1/10 to 10.

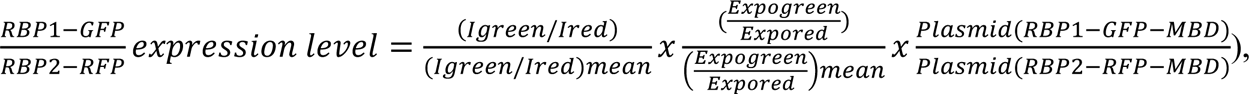

For truncated forms, cells were transfected with a 1:1 ratio.

#### Analysis of images, compartmentalization detection and measurement of determination coefficient

Images were cropped for each cell and intensity and contrast parameters were adjusted with imageJ software. Then, obtained images for each condition were analyzed with a dedicated program developed with CellProfiler to measure the determination coefficient (described in Supplementary Figure S4). Briefly, algorithm detects cells based on their nuclei and then, detects tubular structures (5 to 50 pixels) over a threshold based on their fluorescence intensities. Upper quartile fluorescence intensities for both colors were extracted within the selected structures. Generated data were filtered based on their eccentricity (> 0.9) and background was removed. For each cell a determination coefficient R² was estimated by linear regression between green and red fluorescence intensities from at least 100 values per cell (Microsoft Excel).

#### PLA: Proximity ligation assay

Duolink PLA technology kit (Sigma Aldrich) was used according to the manufacturer’s protocol. HeLa cells were prepared in 96-well plates at 20 000 cells per well concentration. Cells were fixed with PFA 4 % for 20 min at 37°C. Then, cells were incubated in blocking buffer (3 % BSA, 0.1 % triton X-100 in PBS) for 1 h at 37 °C. Primary antibodies for FUS (α-FUS rabbit, mAB ABnova) and TDP-43 (α-TDP-43 mousse, pAB Novus Bio) were diluted to 1:1000 in blocking buffer and incubated for 1 h 30 at room temperature in blocking buffer in humidity chamber. Four replicates from two independent experiments have been done for this condition. U2AF antibodies (Figure 1D) have been used for control (U2AF65 rabbit polyclonal Ab, A303*-*667A, Bethyl, and U2AF65 mouse monoclonal Ab, clone MC3). In parallel, two negative controls were done without primary antibodies or only FUS antibody (Supplementary Figure S3). PLA probes (Minus and Plus, α-rb, α-ms) were diluted 1:5 in the appropriate buffer from the kit, then incubated 1h at 37°C on samples. Cells were washed several times with washing buffer (10 mM Tris, 150 mM NaCl, 0.05 % Tween) before ligation step with ligase (diluted 1:5 ligation stock and 1:40 ligase) incubated 30 min at 37 °C. After washing with PBS, amplification mix was added (diluted 1:50 amplification stock and 1:80 Polymerase) and incubated 100 min at 37 °C. Cells were finally washed with 1X, then 0,1X Tris buffer (200 mM Tris, 100 mM NaCl). DAPI were used to stain nuclei. Samples were stored in PBS without mounting at 4 °C. Acquisitions were done using Opera Phenix Plus confocal microscope with 20x magnification in air using Harmony software. Data were treated with ImageJ and cellProfiler softwares. Each point plotted on the graph represents the percentage of cells with at least one PLA signal (red dot) in the nucleus among 30 cells analyzed.

**Figure 1:**
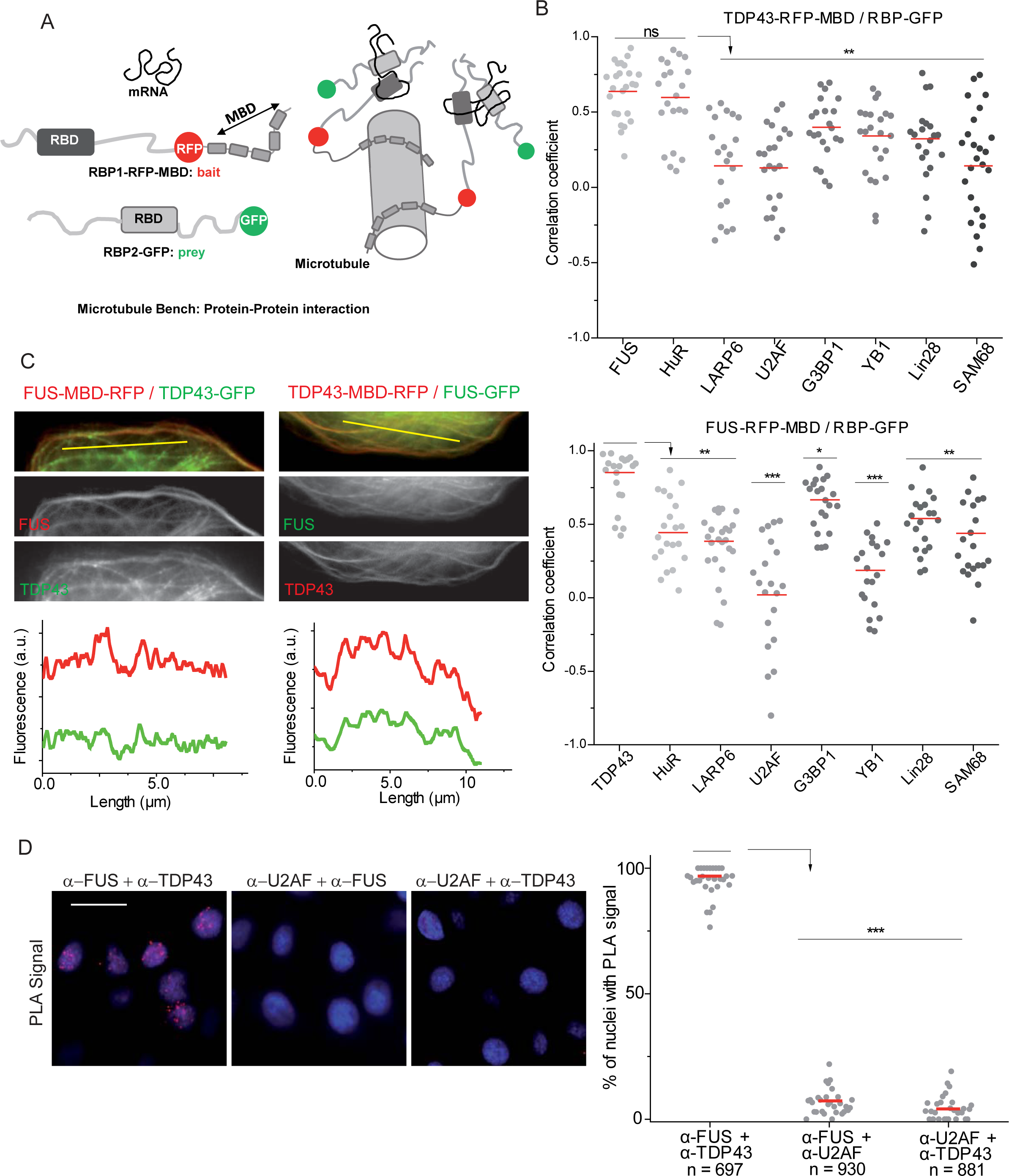
TDP-43 and FUS interact in cellular context. A) Schematic representation of the microtubule bench assay principle. In brief, a protein of interest (bait) fused to microtubule-associated domains of Tau (MBD) and RFP (or GFP) is brought onto microtubules in living cells whereas the presence of a GFP-fused (or RFP) protein partner (prey) on microtubules reveals the interaction by co-localization of the fluorescence signals on microtubule. (B) Scatter plot representing the co-localization level of MBD-fused TDP-43 (upper panel) or MBD-fused FUS (lower panel) with one of the eight tested RBPs. Each data point represents a correlation coefficient between fluorescence intensities from red and green channels along a line crossing microtubules. The plot shows the data from two independent experiments. Red lines show mean values. Significances between correlation coefficients were obtained using t test; *, p < 0.05; **, p < 0.01; ***, p < 0.005; ns, not significant. (C) U2OS cells were co-transfected with the constructions encoding FUS (left panel) or TDP-43 (right panel) fused to RFP-MBD and the plasmid expressing full length TDP-43 (left panel) or FUS (right panel) fused to GFP. The fluorescence intensity from the two channels along the yellow lines are shown below to the respective microphotographs. (D) Left handed panel: Representative images for proximity ligation assay (PLA) revealing the co-localization of TDP-43 and FUS in HeLa cells. Cells were fixed, incubated with anti-FUS, anti U2AF65 or anti-TDP-43 antibodies and with nucleotides probes, ligase and polymerase as described in the Materials and Methods section. Scale bar: 20 µm. Right handed panel: scatter plot representing the percentage of nuclei with a PLA signal. Each point corresponds to 20 to 40 cells analyzed. n: number of cells analyzed. Red lines show mean values. ***, p < 0.005; t-test.

#### Stress granule experiments

Stress granules experiments were performed as previously described [38, 43]. HeLa cells were cultured in DMEM supplemented with 10% FBS in the presence of penicillin and streptomycin (100 μg/mL) (GIBCO Life Technologies). Cell cultures were maintained at 37°C and 5% of CO2 in an incubator. HeLa Cells were plated in 96 well plate (Perkin Elmer). The plasmids were constructed to express, in mammalian cells, FUS and HUR proteins with a GFP tag and TDP-43 wild type or mutants, FUS, HUR, G3BP1, and YB1 bearing an HA tag peptide on N-terminus. For transfection experiments, cells were incubated in presence of 0.3 µg of the plasmid followed by the addition of lipofectamine 2000 reagent (0.2 µL/sample).

Oxidative stress: HeLa cells were treated with puromycin (2.5 µg/ml) and hydrogen peroxide (H2O2) (300 µM) during 90 min at 37°C in a CO2 controlled chamber. After treatment, cells were washed twice with warm-PBS and fixed with 4% paraformaldehyde (PFA) diluted in PBS for 20 min at 37°C. Cells were then incubated with 70% ethanol during 10 min at room temperature followed by incubation in presence of 1 M Tris–HCl pH 8.0 for 5 min.

In situ hybridization: To visualize mRNA, HeLa cells were incubated with a poly-dT oligonucleotide coupled with Cy-2 (Molecular Probes Life Tech.) for 1 hr at 37°C. Washings were carried out using 4× and then 2× SSC buffer (1.75% NaCl and 0.88% sodium citrate, pH 7.0). To visualize HA-tagged protein, cells were incubated overnight at 4°C with an anti-HA mouse, primary antibody (Sigma-Aldrich) diluted (10–3) in blocking buffer containing 0.1% Triton X-100. After washings with PBS, cells were incubated with a secondary goat anti-rabbit IgG antibody (10–3) coupled to Alexa Fluor Plus 594 (Molecular Probes Life Tech.) for 60 min at room temperature. For nuclei visualization, cells were incubated 1 min with DAPI (0.66 mg/mL) (Sigma-Aldrich).

Quantifications were performed with Opera Phenix® Plus High Content Screening System (PerkinElmer) in confocal mode. The Harmony v4.8 software was used to detect automatically the stress granules in cell. The overall cytoplasmic expression of proteins was measured as well as their signal intensity in stress granules, their ratio gives an enrichment in SGs.

#### RNA interference

For the silencing of FUS, HeLa cells were then transfected with 0.15 µg/well of small-interfering RNA (siRNA) duplexes using lipofectamine 2000. A non targeting sequence siRNA (AllStars Negative Control QIAGEN Cat # 10272281) was used as negative control. Cells were transfected with the siRNA and placed in incubator at 37°C for 24 h, then cells were transfected with TDP-25-RFP expression plasmid (0.25 µg/well) and incubated for 24 h at 37°C. Cells were then fixed, incubated with a poly-dT oligonucleotide coupled with Cy-2 and a murin anti-FUS primary antibody (Sigma Aldrich) as previously described.

### *In vitro* experiments

#### In vitro RNA transcription

RNA was produced by *in vitro* transcription procedure as previously described [42]. For this purpose, linearized plasmid pSP72-2Luc, containing two full-length cDNAs encoding *Renilla reinformis* and *Photinus pyralis* luciferases, was used as a template for synthesis of 2Luc mRNA (*∼*3000 nt). HiScribe T7 High Yield RNA Synthesis Kit (NEB) was used for *in vitro* transcription, according to the manufacturer’s protocol. Synthesized RNA was purified using phenol/chloroform extraction.

#### Plasmid preparation for in vitro production of protein

For recombinant protein production, plasmids were produced for full length (FL) and truncated TDP-43 (TDP-25 and TDP-RRM) and FUS thanks to dedicated primers (supplementary table 3). Intermediary plasmids pMS2 were used to add HA tag to the sequence. Fragments were ligated between Nhe1 and BamH1 restriction sites of the plasmid. A second cloning was done to recover the sequence of interest with the HA tag. Finally, the sequence containing the HA tag were transferred in a bacterial expression vector pET22b (Novagen) between Xho1 and HindIII restriction sites. pET22b vector contains a His6 tag for purification, a Lac sequence for protein induction production with IPTG, and an ampicillin gene resistance.

#### Production and purification of proteins

N-terminal His6-tagged recombinant proteins (FL FUS, FL TDP-43 and truncated forms) were expressed in bacteria *Escherichia Coli* BL21 (DE3) strain. After transformation (heat shock 42°C), one bacterial colony grown on LB-agar medium were picked, and transferred in 100 mL 2YT medium with ampicillin 1:1000. This culture was incubated overnight at 37°C under stirring. Then the total volume of preculture was transferred in 1L culture medium and grown until the optical density reached 0.6. Gene expression was induced by addition of 1 mM IPTG and bacteria were grown further for 4 hrs at 37 °C before being collected by centrifugation. Pellet was re-suspended in Tris buffer containing urea (Tris 25 mM, Urea 8 M, NaCl 2 M, 5 mM Imidazole, 0.5 mM DTT, pH 7.4) with PMSF (Sigma) and protease inhibitors (*EDTA-free protease inhibitor cocktail Roche*). Urea is important to keep proteins soluble as they are highly prone to aggregation. Bacterial cells were lysed by sonication (Bioblock Vibracell sonicar, model 72412). Supernatant was finally recovered by ultracentrifugation (75 000g, 40 min, 4°C).

For purification, supernatant was incubated with Ni-Nta-agarose beads (Qiagen) for at least 2 hrs at 4 °C and then loaded in a column equilibrated with Tris-Urea buffer. Several washes with gradient of imidazole (between 10 mM to 100 mM) were done before eluting purified proteins with 250 mM, 500 mM, and 1 M imidazole. All the fractions (washes and elutions) were loaded on a 12% acrylamide gel (SDS-PAGE) for control and to identify fractions containing proteins of interest. Identified fractions were pulled and loaded in desalting column (PD10 Sephadex G-25M, GE-Healthcare) equilibrated with urea buffer (Urea 8 M, Tris 25 mM, NaCl 200 mM, DTT 0.5 mM, pH 7.5) to remove imidazole. Finally, protein samples were concentrated by centrifugation (SpinX^R^ Concentrator 5KMCO or 10KMWCO, Corning). Samples were flash frozen and stored at -80 °C.

#### Electrophoretic mobility shift assay (EMSA)

2Luc mRNA was heated to 80 °C for 1 min then cooled at room temperature. 2Luc mRNA was diluted to obtain 150 ng/well. For the figure 5E, RNA was mixed with proteins in a Hepes buffer (Hepes 20 mM, KCl 25 mM, MgCl2 2m M, DTT 1 mM, pH 7.4) and incubated 15 min at room temperature. For Supplementary Figure 7, proteins were incubated for 50 min alone in Hepes buffer and then 2Luc RNA was added (115 ng/well) for 10 minutes. Samples were loaded on 1 % agarose gel (with BET 5 %, in TAE 1X). Samples were migrated in a TAE 1X buffer under 25 V for 1 hr.

#### Immunofluorescence observation of RBP assemblies in vitro

Proteins were diluted and mixed (with or without 2Luc RNA) in a Tris buffer (Tris 20 mM, KCl 20 mM, MgCl2 2 mM, DTT 1 mM, pH 7.5) and incubated 2 hrs at 37 °C before fixation. Glass slides were pre-treated with poly-lysine 0.2 mg/mL for 15 min, then with glutaraldehyde (2 % in water) for 40 min. After different incubation time, protein samples were deposited on the slides. After 5 min, glutaraldehyde 2 % was added for 15 min to fix samples. Slides were washed in PBS, then NaBH4 (1 mg/mL, freshly prepared solution) was added to bleach glutaraldehyde fluorescence. For immuno-marking, specific primary antibodies against proteins or tags (Supplementary table 4) were used, after a blocking step (BSA 1 %, triton 0.25 % in PBS). During the blocking step, sample could be incubated also for 1h with oligo-dT-dig probe (1:1000, polyT-dig Sigma HA09131354-004). Antibodies were diluted 1:500 in blocking buffer, and incubated 2 hrs at 37°C in humidity chamber. Then secondary antibodies were used to stain the samples. After 2 hrs incubation, slides were washed with PBS and mounted (DAKO^R^ Fluorescent Mounting Medium S3023). Slides were stored at 4 °C. Observations were done with an optical fluorescence microscope (Leica Microsystems DM4B, Hamamatsu C11440 Digital Camera) with an oil immersed objective with 1,6×63× magnification and acquisitions were done with Las X software (2.0.0.14332). Images were treated with ImageJ software. Some acquisitions were obtained using a confocal microscope (TCS SP8, Leica), thanks to imaging platform ImCy (Généthon).

#### Determination of correlation between signals in aggregates

The determination of the correlation between fluorescent signals was done by using an algorithm developed with CellProfiler software. Briefly, the program detects aggregates with an area ranging from 20 and 2000 pixels based on the fluorescence contrast with background (Threshold strategy Global with Ostu method) for green and red channels. A layer is defined for each object to detect clusters (of 2 to 150 pixels) in these aggregates for the two channels. This step is equivalent to form a grid on aggregates in which fluorescence intensity is measured (MeasureObjectIntensity function on CellProfiler software). Then generated data are exported and used to calculate a correlation coefficient (Pearson, with CORREL function of Microsoft Excel software), reflecting the distribution of green and red fluorescence signals in aggregates. This coefficient is comprised between -1 and 1: signals are distinct under 0, heterogeneous around 0, and well mixed close to 1. 8 to 12 images were analyzed for each condition. Area or circularity of aggregates were obtained using MeasureObjectSizeShape function on CellProfiler software. Significance between correlation coefficients were obtained using t test; *, p < 0.05; **, p < 0.01; ***, p < 0.005; ns, not significant

#### AFM imaging

The observation of protein/RNA complexes using AFM was performed as described previously [40]. Full length or truncated proteins were incubated at room temperature alone, mixed or with RNA in a Tris buffer (Tris 10 mM, KCl 15 mM, MgCl2 2 mM, DTT 1 mM, Putrescine 10 mM, pH 6.8). A 10 µL droplet was deposited on freshly cleaved mica surface, and then the surface was quickly immersed in a diluted uranyle acetate solution (0.02 % in water) to fix the sample. Then, the sample was dried with filter paper before imaging. AFM scans were obtained using PeakForce tapping mode in air with Nanoscope V Multimode 8 software (Bruker, Santa Barbara, CA). This model enables continuous force-distance curves recording using Scanasyst-Air probes (Bruker). Images were captured at 1512×1512 pixels at a line rate of 1.5Hz. The “particle analysis” tool on the Nanoscope Analysis software (version 1.70) was used to determine the molecular dimensions of particles (proteins aggregates and mRNA:protein complexes) from at least two independent samples. Basically, a threshold of 2 nm was applied to discard small particles or patterns (uranyl acetate background) and free mRNA from the analysis (supplementary figure S7). The proportion of RBP/RNA complexes is estimated from at least 6 scanned areas (total area = 100 µm²) and ± SD represents the discrepancy between each scanned areas. Significance of areas and ratios were obtained using t test; *, p < 0.05; **, p < 0.01; ***, p < 0.005; ns, not significant

## Results

### Among a set of RBPs, TDP-43 and FUS have the best colocalization score

To determine whether two proteins interact with each other in the cytoplasm, it is possible to use the microtubule network as an intracellular bench [41]. A fusion protein containing an RFP tag and a microtubule-binding domain is overexpressed in cells. It will serve as a bait protein. In parallel, the cells also overexpress a prey protein with a GFP tag (Figure 1A). If the prey protein co-localizes with the microtubule network, it indicates that the two proteins interact directly or indirectly (Supplementary Figure S1). One of the strengths of this analysis is that it reveals interactions in a cellular context, without cell lysis and any purification step. Interaction between TDP-43 and FUS has been compared with a selection of RBPs based on their localization (nucleus/cytoplasm), presence of PrLD, RNA binding motif and involvement in different RNA functions associated to the mRNA life cycle (splicing, translation, stress granule assembly…). When using TDP-43 as a bait, a significant co-localization is mainly detected with two RBPs, FUS and HuR (Figure 1B). The recruitment on the microtubules of the 6 other RBPs tested is significantly less important than that of FUS or HuR (Figure 1C and supplementary Figure 2). When the roles were inverted with FUS used as the bait, TDP-43 had the best co-localization score with FUS among all the RBPs tested as preys. Note that the co-localization score with RBPs used as preys changes depending on whether FUS or TDP-43 was used as bait. Thus, when FUS is the bait protein, G3BP1 is better localized to microtubules than HuR. Moreover, even if some of the RBPs tested as preys have rather a nuclear localization (i.e. FUS, TDP-43, SAM68 or U2AF65), it is still possible to detect their presence on the microtubules. Finally, no prey is detected on the microtubules in the absence of bait, as previously observed [44]. Thus, the co-localization score is systematically the highest with the TDP-43/FUS couple, which reveals a strong affinity between these two RBPs among those tested. To confirm the FUS and TDP-43 co-localization independently of their cytoplasmic localization, proximity ligation assays (PLA) were performed in HeLa cells (Figure 1D and Supplementary Figure S3). The PLA signal indicates a higher co-localization score than previously reported [27] between TDP-43 and FUS in nuclei of Hela cells with nearly 100% of nuclei displaying a signal. Thus, the experimental data carried out in a cellular context confirm the proteomics information from the literature [27, 28, 45] and allow us to conclude on a particular link between TDP-43 and FUS.

### FUS accepts TDP-43 in its compartment

The microtubule network can also be used to determine the ability of proteins to mix or to form their own compartment [39]. To this end, both proteins were fused to a microtubule-binding domain. After their expression in cells, the fusion proteins were brought onto microtubules to generate compartments (demixing) or, on the contrary, a homogeneous phase (mixing, Figure 2A and Supplementary Figure S4) [39]. In the case of RBPs, the formation of compartments involves protein-protein interactions, in particular when they harbor domains of low complexity, but also RNA-protein interactions where RNA provide a scaffold for higher protein assemblies and finally mRNA base pairing. As with polymers, the ability of macromolecules to mix depends on their respective proportion [46]. In order to analyze the formation of compartments when TDP-43 and FUS share the same space, we varied the level of expression of these proteins by modulating the concentration of plasmids during cell transfection. When FUS is highly overexpressed compared to TDP-43, TDP-43 is incorporated within the FUS-rich compartments, as revealed on cell images by yellow microtubules and by an elevated R² value close to 1 (Figure 2C). It corresponds to a limited occurrence of having distinct FUS or TDP-43 rich compartments along microtubules (R² is the square of the correlation coefficient, see Materials and Methods for details). Conversely, when TDP-43 is more expressed than FUS, the microtubules have a less uniform color with successions of red and green clusters indicating the formation of TDP-43- or FUS-rich compartments. Under these conditions, the miscibility between the two analyzed proteins is affected which results in a progressive decrease in R² as the level of expression of TDP-43 increases. There is therefore an asymmetry in the miscibility between the two RBPs.

**Figure 2:**
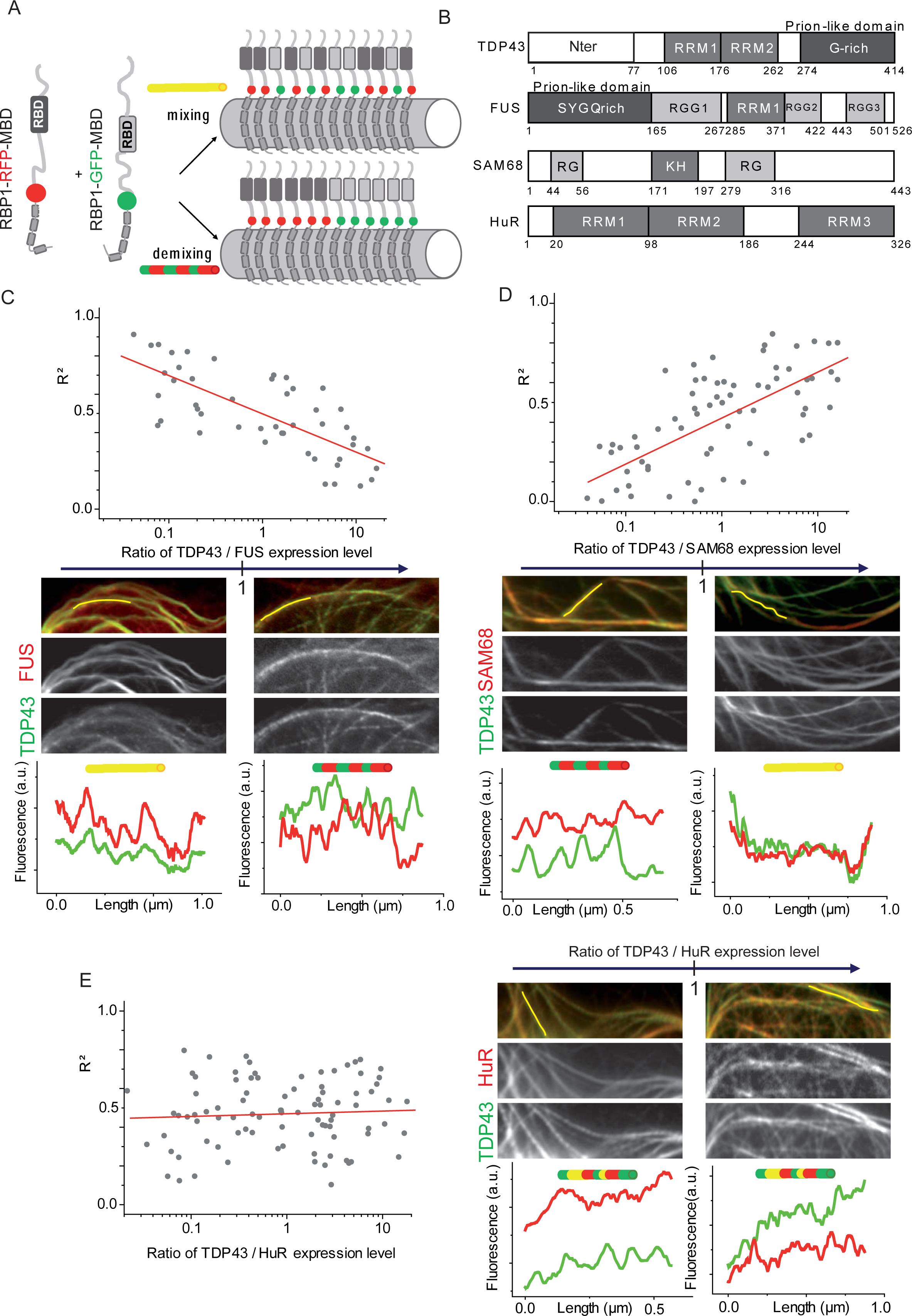
TDP-43 is miscible in FUS-rich compartments in cellular context. (A) Two RBPs as indicated are confined on the microtubule network (fused to RFP/GFP-MBD) to visualize their mixing/demixing in U2OS cells. Mixing: yellow microtubules. Demixing: red and green microtubules. (B) Schematic representation of the domains of RNA-binding proteins used in the microtubule bench assay to reveal their mixing according to their expression level. (C) Upper panel: scatter plot representing the mixing between TDP-43 and FUS, both fused to MBD according to the expression level of each construct. Each data point represents a value of determination coefficient R² calculated for one cell as described in the Materials and Methods section. Lower panel: representative images for a low (left handed panel) and a high (right handed panel) TDP-43/FUS expression level ratio. The fluorescence intensity from the two channels along the yellow lines are shown below to the respective microphotographs. (D) Same as (C) with the mixing between TDP-43 and SAM68 fused to MBD. (E) Same as (C) with the mixing between TDP-43 and HuR fused to MBD.

Then we checked whether TDP-43 has the same exclusion behavior with other RBPs or whether it is specific to its association with FUS. Conversely, as the compartments of FUS seem to accept the presence of TDP-43, we wanted to probe whether this behavior is also valid for other RBPs. To address these points, we selected two RBPs that have different domains and aggregation propensity. SAM68 (SRC associated in mitosis of 68-kDa) is a protein that harbors several low complexity domains (RG rich domains) and an RNA-binding domain (KH domain) (Figure 2B) with a strong propensity for multimerization or even aggregation: SAM68 is the main component of SAM68 nuclear bodies (SNBs) [42, 47–49]. Either when SAM68 is brought together with TDP-43 or FUS on microtubules, the formation of distinct SAM-68 rich compartments is clearly evidenced when the expression level of SAM68 is high (Figure 2D and Supplementary Figure S5). The ability of SAM68 to multimerize is certainly involved in its strong propensity to form compartments on its own. Unlike SAM68, HuR (Human antigen R) does not harbor LCDs but rather multiple RRMs that can form multimers [50]. Whatever the overexpression level of HuR and the nature of the other RBP partner present along microtubules (TDP-43 or FUS), R² remains relatively stable and in a range of values indicating an average miscibility between HuR and TDP-43 (or FUS) (Figure 2E and Supplementary Figure S5). In summary, TDP-43-rich compartments exclude FUS while, on the contrary, FUS-rich compartments accept TDP-43 in its environment.

### *In vitro*, TDP-43 and FUS form structurally different assemblies

To understand the difference in the ability of a protein to accept the other in its environment, TDP-43 and FUS were purified and incubated separately before being fixed on a glass slide for their observation by fluorescence optical microscopy (Figure 3A and B). This process allows the structural characterization of the higher order assemblies of TDP-43 and FUS. FUS forms assemblies whose size is stable over time (Figure 3C) and with a circular shape (Figure 3D and supplementary Figure S6). There is a clear structural resemblance between these assemblies and the droplets of FUS detected during the Liquid-Liquid Phase Separation (LLPS) process [51–57]. TDP-43 forms different assemblies than FUS (Figure 3B), increasing in size with incubation time (Figure 3C) or concentration (supplementary Figure S6) and exhibiting a filamentous appearance (Figure 3D and supplementary video SV1). These two proteins were then incubated in the same buffer for 2 hrs and in variable proportion (Figure 3E). By gradually increasing the fraction of TDP-43, we move from structures having a morphology close to FUS to that close to TDP-43. As soon as a small fraction of TDP-43 is added, we evidence an association with the FUS droplets leading to the formation of clusters associated with a reduction in circularity (Figure 3F). However, the high correlation coefficient indicates a miscibility between TDP-43 and FUS in these FUS-rich clusters. When FUS is incubated with a higher fraction of TDP-43 (ie FUS/TDP-43 molar ratio of 1/3 and beyond), the structure of the assemblies evolves to reach the ones observed in Figure 3B for TDP-43 alone. The results also indicate a significant decrease in the circularity of the assemblies, whether the fluorescence signal coming from FUS or TDP-43 was used to estimate the circularity (Figure 3F and G respectively). Finally, at a FUS/TDP-43 molar ratio of 3/1, the correlation coefficient collapses, and corresponds to the spatial separation between TDP-43 and FUS with the formation of TDP-43-rich or FUS-rich structures (Figure 3H). Thus, FUS only seems able to solubilize TDP-43 when present at low concentration, preventing the formation of distinct TDP-43-driven structures, which occurs at higher TDP-43 concentrations inside FUS-rich compartments. In the latter case, we can qualify the TDP-43-rich assemblies still interacting with the FUS compartment as subcompartments.

**Figure 3:**
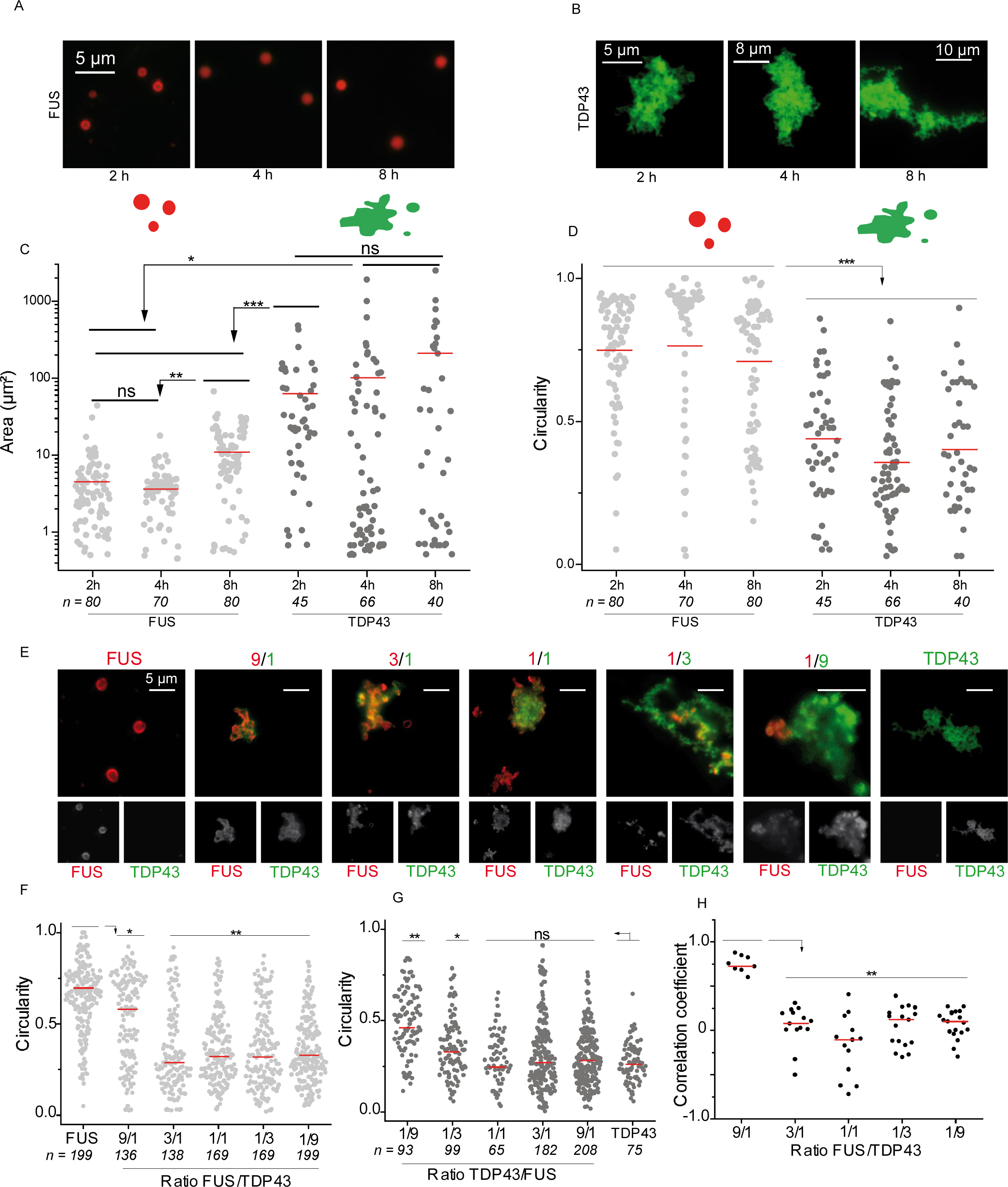
*In vitro*, FUS limits the segregation of TDP-43. (A) Images of FUS assemblies revealed by immunofluorescence after incubation of FUS proteins at 5 µM for different incubation times. Scale bar: 5 µm. (B) Images of TDP-43 assemblies revealed by immunofluorescence after incubation of TDP-43 proteins at 5 µM for different incubation times. (C) Scatter plot representing the area of FUS and TDP-43 assemblies along the incubation time. The plot shows the data from two independent experiments. n: number of assemblies analyzed. Red lines show mean values. Significances between areas of FUS and TDP-43 assemblies were obtained using t test; *, p < 0.05; **, p < 0.01, ***, p < 0.005, ns, not significant. (D) Scatter plot representing the circularity of FUS and TDP-43 assemblies along the incubation time. The plot shows the data from two independent experiments. n: number of assemblies analyzed. Red lines show mean values. Significances between circularity of FUS and TDP-43 assemblies were obtained using t test; ***, p < 0.005. (E) Representative images of FUS/TDP-43 assemblies after incubation of the two proteins for two hours at different ratios ranging from 9 to 1/9. Top: merged signal from FUS-RFP and TDP-43 GFP. Bottom: images of the same assemblies but from FUS-RFP (left) or TDP-43-GFP (right). Scale bar: 5 µm. (F) Scatter plot representing the circularity of FUS rich phases in FUS/TDP-43 assemblies according to the FUS/TDP-43 ratio. Only the fluorescence signal from FUS-RFP was analyzed. The plot gathers the data from two independent experiments. n: number of assemblies analyzed. Red lines show mean values. Significances between circularity of FUS/TDP-43 assemblies were obtained using t test; *, p < 0.05; **, p < 0.01. (G) Scatter plot representing the circularity of TDP-43 rich phases in FUS/TDP-43 assemblies according to the FUS/TDP-43 ratio. Only the fluorescence signal from TDP-43-GFP was analyzed. The plot gathers the data from two independent experiments. n: number of assemblies analyzed. Red lines show mean values. Significances between circularity of FUS/TDP-43 assemblies according to the protein ratio were obtained using t test; *, p < 0.05; **, p < 0.01; ns, not significant. (H) Scatter plot representing the correlation coefficient between the fluorescence signals of FUS-RFP and TDP-43-GFP. Each data point represents a correlation coefficient between fluorescence intensities from red and green channels in aggregates with an area comprised between 20 and 2000 pixels. The plot shows the data from two independent experiments. Lines show mean values. Significances between correlation coefficients were obtained using t test; **, p < 0.01.

### *In vitro*, RNA improves the miscibility of these two proteins

The assemblies of the TDP-43 and FUS are modulated by the presence of RNA. Those of FUS remain spherical in the presence of RNA but their size decreases when the FUS/RNA molar ratio decreases (Figure 4A). For TDP-43, the influence of RNA on the formation of its higher order assemblies is more visible. For a protein/RNA ratio of 10/1, RNA favors the formation of large assemblies characterized by a significant increase in the average area. At lower ratio, large TDP-43 assemblies dissociate, as expected owing the buffering action of RNA (Figure 4B) [8]. Then, we probed whether the structures of TDP-43 in interaction with FUS depends on RNA. RNA was first incubated in the presence of FUS for few minutes before TDP-43 was added in variable proportion until becoming dominant while the protein/RNA ratio was fixed at 1/10 (Figure 4C). The structures observed are quite similar to those obtained in Figure 3E without RNA (Supplementary Video SV2 and SV3). The typical pattern of FUS (spherical assemblies that associate in the form of clusters) is gradually replaced by that of TDP 43 with more massive assemblies of filamentous appearance. When the proportion of TDP-43 increases, the correlation coefficient decreases but the starting point of the demixing process is shifted towards higher proportions of TDP-43 compared to what was observed in the absence of RNA (Figure 4E). During the opposite process, i.e. when TDP-43 is first incubated with RNA before FUS is added with increasing proportions, we remarked that FUS is hardly incorporated into the assemblies of TDP-43 as evidenced by a low correlation coefficient (Figure 4D and F). The FUS/TDP-43 ratio should be higher than 1 to trigger the mixing of the two proteins, characterized by a gradual increase of the correlation coefficient (Figure 4F). We then wondered whether the concentration of RNA also modulate the ability of FUS and TDP-43 to mix. When the amount of RNA increases, i.e. the molar ratio Protein/RNA nucleotide decreases from 1 to 0.1, with a constant FUS/TDP-43 molar ratio of 3, the colocalization between the two proteins clearly increased (Figure 5A).

**Figure 4:**
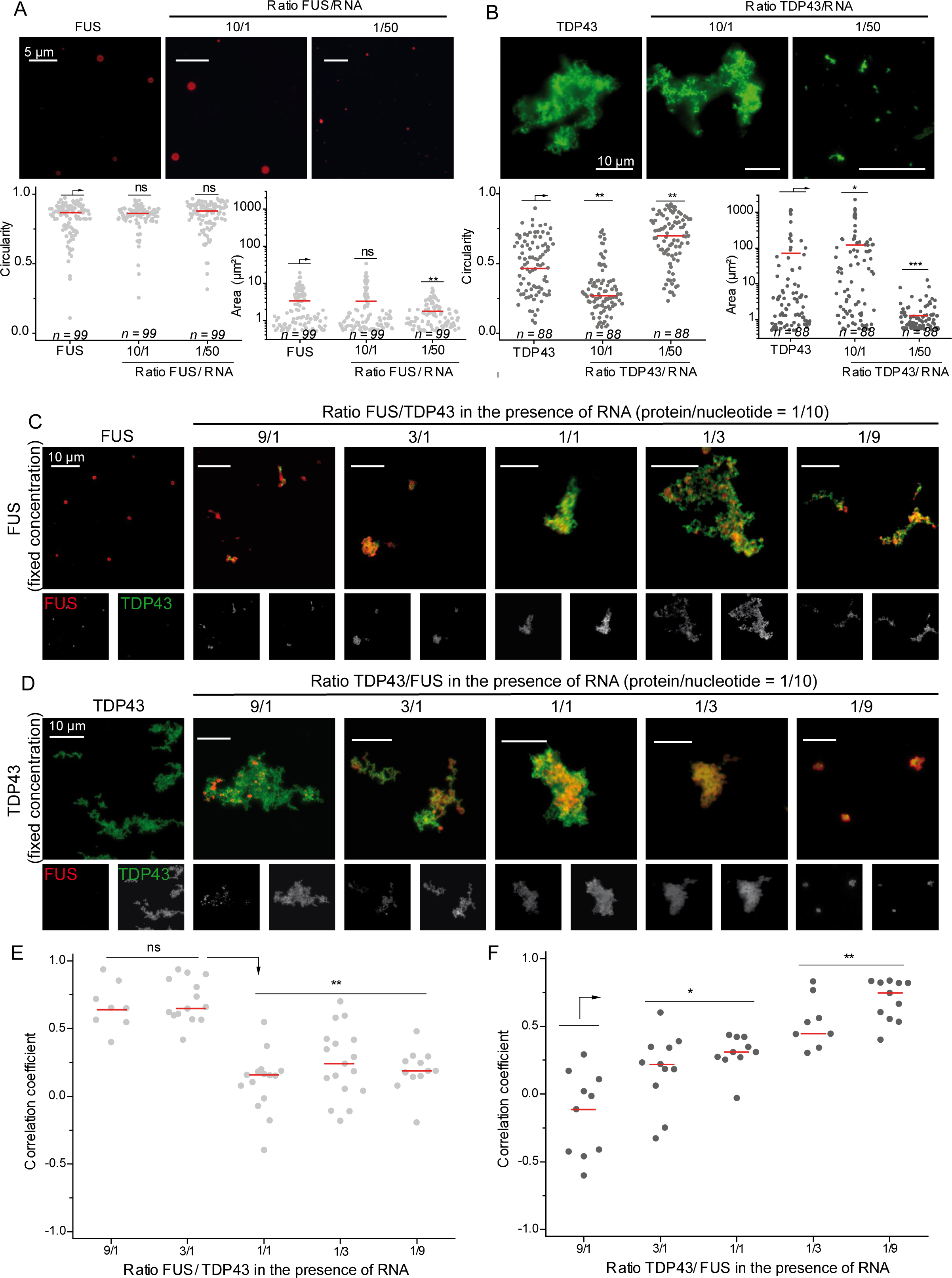
RNA improves the miscibility of TDP-43 in FUS-rich assemblies. (A) Upper panel: images of FUS assemblies revealed by immunofluorescence after incubation of FUS proteins for two hours at 5 µM with or without 2Luc RNA. Lower panel: scatter plot representing the circularity (left) and area (right) of FUS assemblies according to the protein to nucleotide ratio. The plot gathers the data from two independent experiments. n: number of assemblies analyzed. Lines show mean values. Significances between circularity (or area) of FUS assemblies were obtained using t test; **, p < 0.01; ns, not significant. (B) Upper panel: images of TDP-43 assemblies revealed by immunofluorescence after incubation of TDP-43 proteins for two hours at 5 µM with or without 2Luc RNA. Lower panel: scatter plot representing the circularity (left) and area (right) of TDP-43 assemblies according to the protein to nucleotide ratio. The plot gathers the data from two independent experiments. n: number of assemblies analyzed. Red lines show mean values. Significances between circularity (or area) of TDP-43 assemblies were obtained using t test; *, p < 0.05; **, p < 0.01; ***, p < 0.005. (C) Images of FUS/RNA assemblies in the presence of increasing amounts of TDP-43. FUS was incubated two minutes with 2Luc RNA before TDP-43 addition and further incubation for two hours. FUS/TDP-43 ratios were ranging from 9 to 1/9 with a concentration of FUS (5 µM) constant and a fixed protein/RNA nucleotide of 1/10. (D) Images of TDP-43/RNA assemblies in the presence of increasing amounts of FUS. TDP-43 was incubated two minutes with 2Luc RNA before FUS addition and further incubation for two hours. TDP-43/FUS ratios were ranging from 9 to 1/9 with a concentration of TDP-43 (1.67 µM) constant and a fixed protein/RNA nucleotide of 1/10. (E) Scatter plot representing the correlation coefficient between the fluorescence signals of FUS and TDP-43 in the presence of 2Luc RNA in the conditions described in (C) (left handed panel) on in (D) (right handed panel). Each data point represents a correlation coefficient between fluorescence intensities from red and green channels and only aggregates with an area comprised between 20 and 2000 pixels are selected. The plot shows the data from two independent experiments. Red lines show mean values. Significances between correlation coefficients were obtained using t test; **, p < 0.01, ns, not significant. (F) Scatter plot representing the correlation coefficient between the fluorescence signals of FUS and TDP-43 in the presence of 2Luc RNA in the conditions described in (D). Each data point represents a correlation coefficient between fluorescence intensities from red and green channels and only aggregates with an area comprised between 20 and 2000 pixels are selected. The plot shows the data from two independent experiments. Lines show mean values. Significances between correlation coefficients were obtained using t test; *, p < 0.05; **, p < 0.01.

**Figure 5:**
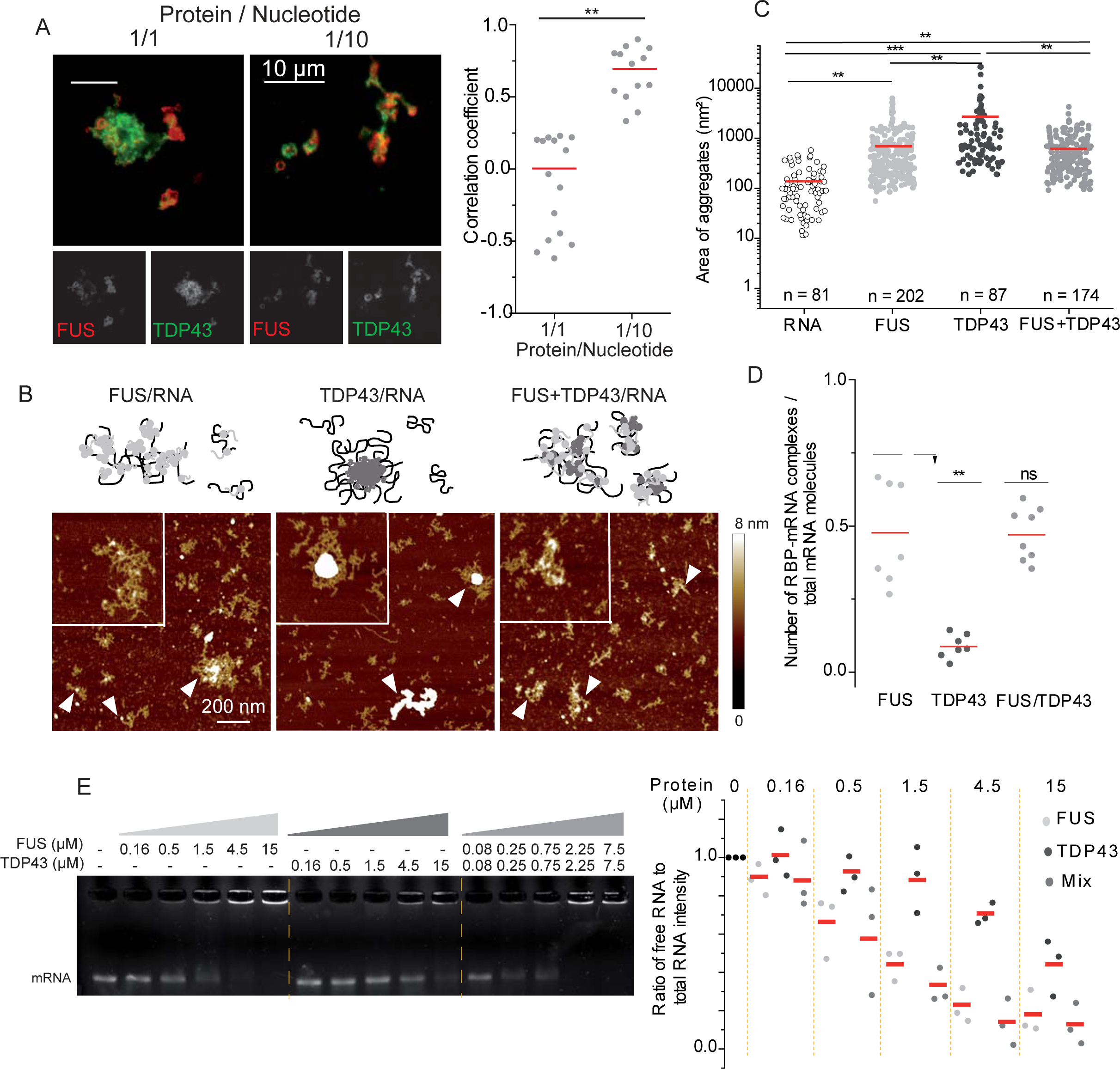
FUS promotes the TDP-43 distribution on 2Luc RNA. (A) Left handed panel: images of FUS/TDP-43 assemblies revealed by immunofluorescence in the presence of various amount of 2Luc RNA. FUS (5 µM) and TDP-43 (1.67 µM) were incubated for two hours with 2Luc RNA with protein/nucleotide ratio of 1/1 or 1/10. Right handed panel: scatter plot representing the correlation coefficient between the fluorescent signals of FUS and TDP-43 according to the protein/nucleotide ratio. The plot shows the data from two independent experiments. Lines show mean values. Significances between correlation coefficients were obtained using t test; **, p < 0.01. (B) Atomic force microscopy images and zooms in on specific assemblies of FUS/RNA, TDP-43/RNA and of an equimolar mixture of the two RBPs with 2Luc RNA. Proteins (500 nM) were incubated with 2Luc RNA (protein/nucleotide ratio of 1/100) for 15 minutes before sample deposition and fixation. White arrows pointed on RNA/RBPs complexes. Z scale 8 nm. (C) Scatter plot representing the area of RBP/RNA assemblies observed on AFM images following the same conditions as in (B). Only RBP/RNA complexes and not free RNA molecules are taken into account using a height threshold of 2 nm. The plot gathers the data from two independent experiments. n: number of assemblies analyzed. Lines show mean values. Significances between areas of RBP/RNA assemblies were obtained using t test; **, p < 0.01, ***, p < 0.005. (D) Scatter plot of the proportion of RBP/RNA complexes compared to total structures (complexes and free RNAs) adsorbed on mica surface and observed on AFM images. Each point corresponds to the same surface analyzed, here 100 µm². The plot gathers the data from two independent experiments. Lines show mean values. Significances between FUS/RNA, TDP-43/RNA and (FUS+TDP-43)/RNA samples were obtained using t test; **, p < 0.01; ns, not significant. (E) RNA mobility shift assay demonstrating the direct interaction between RBPs and RNA. 150 ng of 2Luc RNA were incubated with increasing concentration of TDP-43 (for lane 2 to 6) or FUS (for lane 7 to 11). For TDP-43 + FUS mixes, each protein concentration was divided per 2 to maintain the same global protein/nucleotide ratio with a 1:1 TDP-43/FUS ratio. Quantification of the free RNA band intensity of each lane (normalized, lane 1) on triplicate.

To further explore the consequences of such asymmetric miscibility between FUS and TDP-43 on the structure of protein/RNA complexes, we used high resolution Atomic Force Microscopy (AFM) enabling nanometer scale analysis of multimolecular assemblies. FUS was incubated with RNA for few minutes to analyze the formation of RNA/FUS complexes (protein/nucleotide ratio fixed at 1/100). FUS is quite homogenously distributed among the RNAs (Figure 5B and 5D, Supplementary Figure S7) since approximately half of the RNAs adsorbed on the surface are interacting with one or more FUS proteins, leading to the formation of large assemblies weakly compacted or isolated complexes. For the same incubation time and protein/nucleotide ratio, TDP-43 repartition on RNA is less homogeneous (Figure 5B and D, supplementary Figure S7). TDP-43 tends to accumulate on few RNAs thus forming highly compacted and large structures (arrows in Figure 5B and supplementary Figure S7). When equimolar proportions of TDP-43 and FUS are incubated with RNA for the same incubation time and protein/nucleotide ratio, numerous structural changes are observed in the protein/RNA complexes. First, the proportion of RNAs complexed with proteins is closed to that observed with FUS and RNA alone (Figure 5D). Second, the area of RNA/protein assemblies decreases considerably compared to TDP-43/RNA samples (Figure 5C). In parallel, the large compact structures observed in the TDP-43/RNA samples are no longer detected. Thus, at the micrometric scale (Figure 4C and 4D), the association between TDP-43 and FUS in the presence of RNA seems to favor the formation of large assemblies in which we noticed the presence of clusters. Therefore, at the nanometer scale, the results indicate that the presence of FUS limits the capacity of TDP-43 for massive self-assembly in distinct TDP-43-rich structures to promote the formation of TDP-43 cluster embedded in FUS/RNA assemblies. The buffering action of FUS toward TDP-43 obtained at the single molecule level was further corroborated by gel shift assays. Indeed, the combination of TDP-43 and FUS leads to their association with a slightly higher fraction of RNA than when RNA is incubated with only one of the proteins. Importantly, TDP-43 alone poorly associates with mRNA, which should have considerably reduced the capacity of an equimolar mixture of TDP-43 and FUS to associate with RNA compared to FUS alone as we kept constant the total RBP concentration (Figure 5E). In addition, the presence of FUS within pre-incubated TDP-43 assemblies favors their association with RNA compared to TDP-43 and FUS assemblies considered independently (Supplementary Figure S8). Thus, the interplay between FUS and TDP-43 regulates their mutual higher order assemblies in the presence of RNA to preserve the functional binding of TDP-43 to mRNA.

### TDP-43 and FUS colocalization in stress granules is mediated by TDP-43 cooperative binding to RNA

TDP-43 and FUS are nuclear proteins that can shuttle to the cytoplasm upon stress exposure and assemble into stress granules (SGs) [58, 59]. Here, we overexpressed GFP-RBPs and HA-tagged RBPs in HeLa cells. Then, stress granule assembly was triggered by exposure to a combined puromycin/hydrogen peroxide treatment. Puromycin causes premature chain termination, which facilitates the appearance of SGs in most hydrogen peroxide-treated HeLa cells (Figure 6A). Puromycin/hydrogen peroxide treatment allows to better detect cytoplasmic FUS and TDP-43 recruitment in SGs than with arsenite treatment, mostly because FUS and TDP-43 translocated in the cytoplasm after H2O2 treatment [43]. Cells were imaged using an automatic HCS imager operating in confocal mode at high resolution and the RBP enrichment in stress granules compared to the cytoplasm was measured. Depending on the RBP overexpressed, the relative FUS enrichment in the SGs varies but is higher when TDP-43 is overexpressed (Figure 6B), in line with the miscibility previously observed *in vitro* and on the microtubule network (Figure 1B and 5B). In addition, the TDP-43 enrichment in the SGs is promoted by the overexpression of FUS compared to HuR or GFP alone (Figure 6C). In order to highlight the importance of RNA binding in the ability of FUS and TDP-43 to coexist in the same condensates, we overexpressed FUS or HuR with a TDP-43 mutant (TDP43 G146A) in which the interface between the RRMs of two adjacent TDP-43 bound to RNA is affected, leading to an impaired cooperativity [38] (Figure 6D). It appears that the average mRNA enrichment in SGs is relatively constant independently of the overexpressed protein. In particular, the overexpression of TDP-43 WT or G146A mutant with FUS, do not exhibit significant differences in the mRNA enrichment in SGs (Figure 6E). However, we notice a strong decrease of the enrichment of the TDP-43 G146A in the SGs compare to TDP-43 WT independently of the co-expressed RBPs, here FUS or HuR (Figure 6G). In addition, the enrichment of FUS in SGs, but not HuR, decreases when the cooperativity-defective TDP-43 mutant is overexpressed (Figure 6F). This analysis in a cellular context demonstrates that TDP-43 binding to mRNA is at the heart of the interaction between FUS and TDP-43.

**Figure 6:**
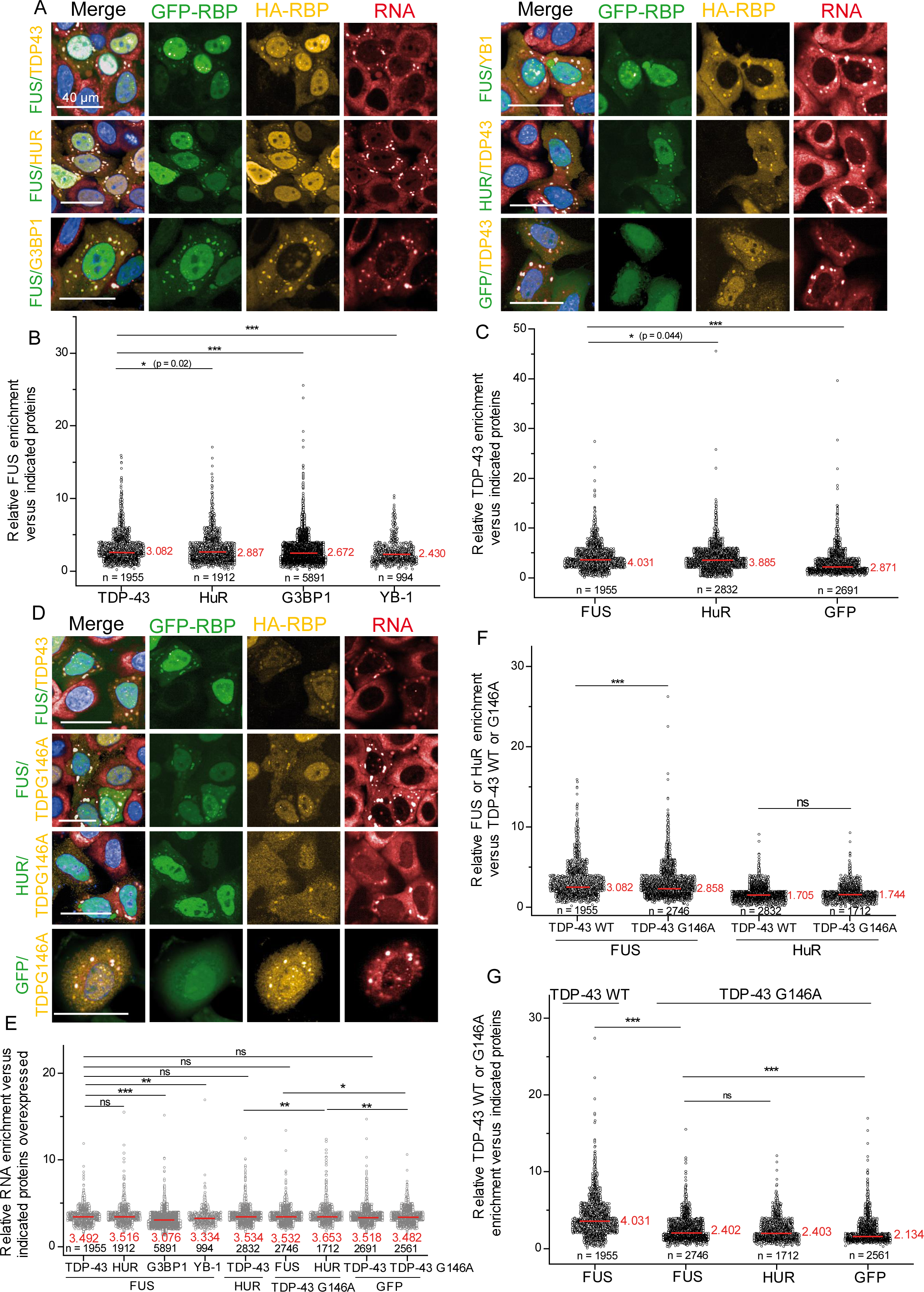
FUS and TDP-43 colocalization in SGs is impaired by a cooperativity-defective mutation in TDP-43. (A) Representative images of HeLa cells overexpressing HA and GFP tagged RBP exposed to H202 and puromycin treatments to trigger SG assembly. Scale bar 40 µm. (B) Scatter plot representing the relative FUS-GFP enrichment in SGs versus overexpressed HA-RBPs. Significances between FUS enrichment levels were obtained using t test; *, p < 0.05; ***, p < 0.005. n: number of cells analyzed; in red mean value. (C) Scatter plot representing the relative TDP-43-HA enrichment in SGs versus overexpressed GFP-RBPs or GFP alone as a control. Significances between TDP-43 enrichment levels were obtained using t test; *, p < 0.05; ***, p < 0.005. (D) Representative images of HeLa cells overexpressing HA and GFP tagged RBP exposed to H202 and puromycin treatments to trigger SG assembly. Scale bar 40 µm. (E) Scatter plot representing the relative RNA enrichment in SGs versus overexpressed HA and GFP tagged RBPs. Significances between RNA enrichment levels were obtained using t test; *, p < 0.05; **, p < 0.01, ***, p < 0.005; ns, not significant. (F) Scatter plot representing the relative FUS-GFP or HuR-GFP enrichment in SGs versus TDP-43 and TDP-43 G146A cooperative-defective mutant. Significances between FUS and HuR enrichment levels were obtained using t test; ***, p < 0.005. (G) Scatter plot representing the relative TDP-43-HA or TDP-43 G146A-HA enrichment in SGs versus overexpressed GFP-RBPs or GFP alone as a control. Significances between WT and mutant TDP-43 enrichment levels were obtained using t test; ***, p < 0.005; ns, not significant.

### FUS is not able to solubilize TDP-25

FUS and TDP-43 are able to bind to the same RNA molecules leading to a higher miscibility of FUS and TDP-43 than in the absence of RNA. To determine whether RNA is the only factor controlling the miscibility of these proteins, the miscibility of truncated proteins was studied using the microtubule bench (Figure 7A and supplementary figure S10). Among the truncations, we selected TDP-25 and TDP-35 which are detected in the frontal and temporal lobes of ALS and FTLD patients [2, 60]. In general, the miscibility of TDP-43 truncation mutants with full length TDP-43 (FL TDP-43) appears to negatively correlate with the length of the truncated parts (Figure 7B). Interestingly, the two RRMs alone are not sufficient to induce a very high miscibility with the FL TDP-43. Both the N-terminal domain known to multimerize [61] and the RRM domains are required in the truncated mutant to maintain a partly preserved miscibility with FL TDP-43 (Figure 7B). When FL TDP-43 interacts along microtubules with truncated FUS, the miscibility with TDP-43 also decreases compared to the FL FUS (Supplementary Figure S10). No truncation of FUS seems to display a preserved miscibility with FL TDP-43: all the domains may contribute to the observed miscibility. If FL FUS interacts along microtubule network with truncated forms of TDP-43, we first notice that the miscibility between FL FUS and TDP-43 RRMs is closed to that of FL TDP-43 (Figure 7C), which highlights the importance of the TDP-43 RRMs for the mixing with FUS. Consistently, the TDP-25 truncation in which the RRM1 has been removed as well as part of the RRM2 do not mix with FL FUS. Thus, it seems that the truncations able to bind to RNA are more prone to associate with FUS compartments than those for which these domains are partially or totally truncated. Moreover, truncated TDP-43 that still contain the PrLD domain have lower miscibility with FL FUS, which indicates that the PrLD domain promotes the spatial segregation of TD-43 or its sub-compartmentalization.

**Figure 7:**
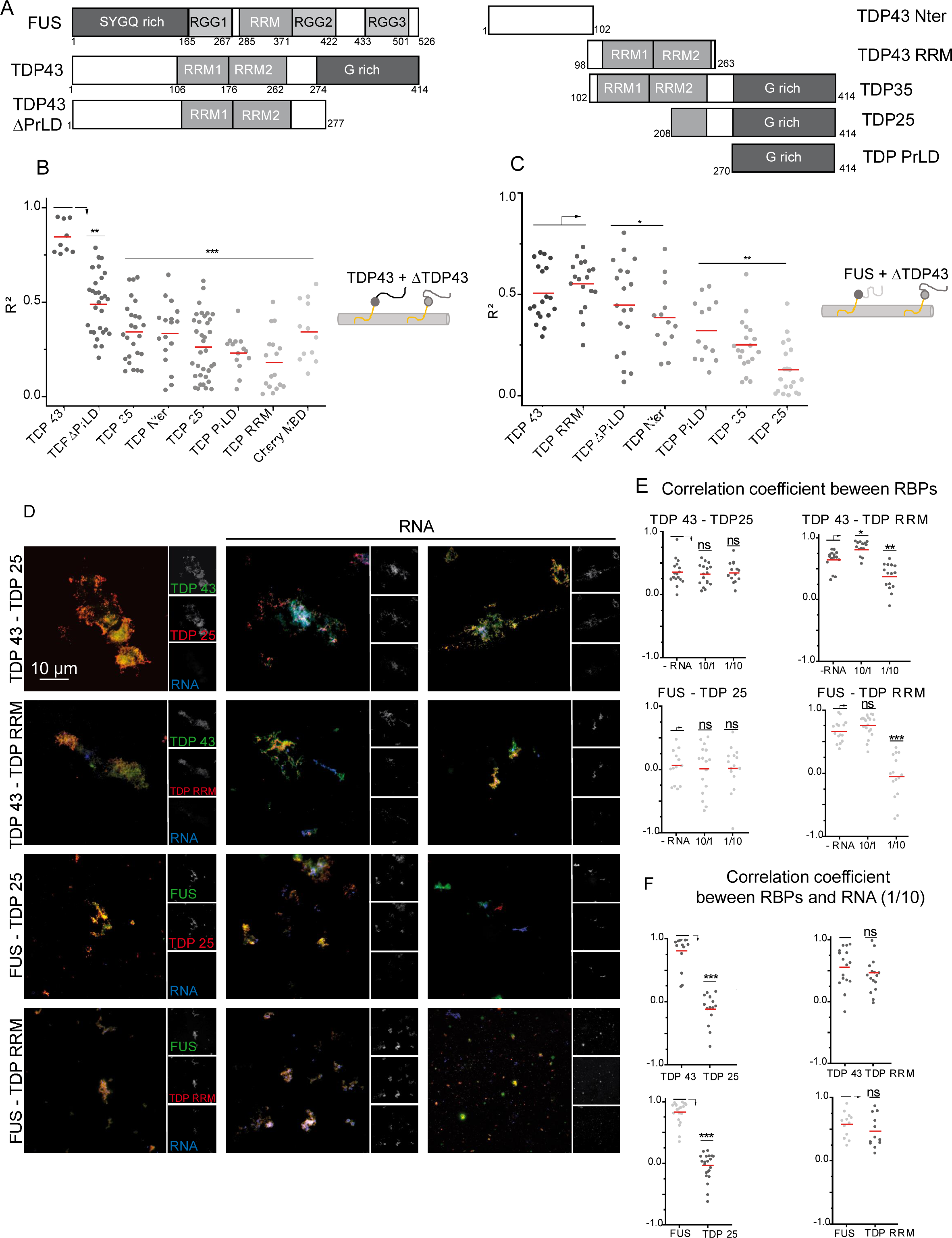
FUS does not mix with TDP-43 truncations with an impaired RNA binding. (A) Schematic representation of the domains of TDP-43 truncations used in the microtubule bench assay to reveal their mixing with full length FUS or full length TDP-43. (B) Scatter plot representing the mixing between full length TDP-43 and TDP-43 truncated forms (ΔTDP-43), both fused to MBD. Each data point represents a value of determination coefficient R² calculated for one cell. Cherry-MBD construct is considered as a control as no interaction (attraction or repulsion) is expected between TDP-43 and Cherry protein. Red lines show mean values. Significances between determination coefficients were obtained using t test; **, p < 0.01; ***, p < 0.005. (C) Scatter plot representing the mixing between FUS full length and TDP-43 truncated forms, both fused to MBD. Each data point represents a value of determination coefficient R² calculated for one cell. Red lines show mean values. Significances between determination coefficients were obtained using t test; *, p < 0.05; **, p < 0.01. (D) Images of assemblies generated by the incubation for 2 hours of an equimolar mix of FUS (or TDP-43) with TDP25 (or TDP RRM) in the absence (first lane) or the presence of 2Luc RNA (in situ hybridization with oligo-d(T) probes). Protein to nucleotide ratio ranging from 10/1 (second lane) to 1/10 (third lane) with a total protein concentration of 5 µM. Full length FUS or TDP-43 were incubated less than one minute with 2Luc RNA before the addition of the equimolar amount of truncated forms of TDP-43. Scale bar: 10 µm. (E) Scatter plot representing the correlation coefficient between the fluorescence signals of FL FUS (or FL TDP-43) and truncated TDP-43. Each data point represents a correlation coefficient between fluorescence intensities from red and green channels in assemblies with an area comprised between 20 and 2000 pixels. The plot shows the data from two independent experiments. Red lines show mean values. Significances between correlation coefficients were obtained using t test; *, p < 0.05; **, p < 0.01, ***, p < 0.005; ns, not significant. (F) Scatter plot representing the correlation coefficient between the fluorescence signals of proteins (FL TDP-43 or FL FUS or truncated TDP-43) and 2Luc mRNA. Each data point represents a correlation coefficient between fluorescence intensities of red or green channels and blue channel in aggregates with an area comprised between 20 and 2000 pixels. The plot shows the data from two independent experiments. Red lines show mean values. Significances between correlation coefficients were obtained using t test; ***, p < 0.005; ns, not significant.

In order to confront the results obtained in cells with *in vitro* analyses, the assemblies formed by incubating truncation mutants with FL TDP-43 or FUS during two hours were observed by fluorescence optical microscopy (Figure 7D). FL TDP-43 and TDP-25 form massive assemblies which are not dissolved upon increasing RNA levels. The correlation between these two proteins remains low and not modulated by the presence of RNA (Figure 7E). Moreover, RNA labeling makes it possible to detect a preferential co-localization between FL TDP-43 and RNA (Figure 7F), which seems consistent with the absence of an efficient RNA-binding domains in TDP-25. Conversely, in the presence of RNA, TDP-43 RRM produces smaller assemblies whether alone or in combination with FL TDP-43 than FL TDP-43 alone. RNA increases the miscibility between TDP-43 and TDP-RRM at moderate concentrations and then decreases the miscibility at high RNA concentration, which is certainly explained by the ability of FL TDP-43 and RRM to bind to RNA. Indeed, RNA has the same co-localization score with the two constructs (Figure 7F). When TDP-43 is replaced by FUS, the size of the assemblies decreases in line with the results obtained for FL FUS alone (Figure 4A). The correlation of the fluorescence signals coming from FUS and TDP-25 is low either in the presence of RNA or not. It indicates that the two proteins poorly mix with each other while that the presence of RNA fails to promote their association. This result is consistent with FUS being the only protein able to bind RNA unlike TDP-25. In the presence of TDP RRM, the correlation between the signals of TDP RRM and FUS is quite high even in the absence of RNA. The mixing between FL FUS and TDP-43 RRM slightly increases in the presence of RNA before decreasing sharply when RNA is in excess, which is similar to the behavior observed when TDP-43 RRM interacts with FL TDP-43.

The set of results that we acquired raises the question of a lack of interactions between FUS and TDP-25 leading to the aggregation propensity of TDP-25. We then explored this point through an atomic force microcopy analysis. We demonstrated previously that FUS favors the dispersion of FL TDP-43 on mRNA by using high resolution imaging (Figure 5B). When FUS is incubated with RNA, FUS interacts homogenously with most of the RNAs (white arrowheads in Figure 8A) before gradually gathering FUS and RNA together in the form of granules, in which the presence of RNA is still visible (like in Figure 5B). Compared to FUS, TDP-25 is found in the form of isolated proteins adsorbed on the surface which can also form compact aggregates where the presence of RNA cannot be detected (orange arrowheads in Figure 8A). When FUS and TDP-25 are incubated at the same time with RNA, some RNA-protein complexes are observed (white arrowheads) with many RNA-free proteins adsorbed on the surface. This system then evolves towards different structural assemblies, one closed to that observed for the FUS/RNA complexes (white arrowheads), the other massive and compact without apparent RNA fragments (orange arrowheads). Thus, FUS is not able to solubilize TDP-25 fragment which could then freely form aggregates independently of the presence of RNA. We confirmed this results in cells exposed to a combined puromycin/hydrogen peroxide treatment (Figure 8B). In these conditions, FUS accumulates in stress granules detected by the mRNA enrichment while TDP-25 is absent from these granules. After reducing the expression of FUS by siRNA (Figure 8C), no significant difference in TDP-25 enrichment in stress granules was detected (Figure 8D). Importantly the mRNA enrichment in the granules is independent on the FUS expression level (Supplementary Figure S11). In addition, we noticed the presence of numerous small TDP-25 rich granules as already observed [62]. These small aggregates are independent of the expression level of FUS and are not enriched in mRNA.

**Figure 8:**
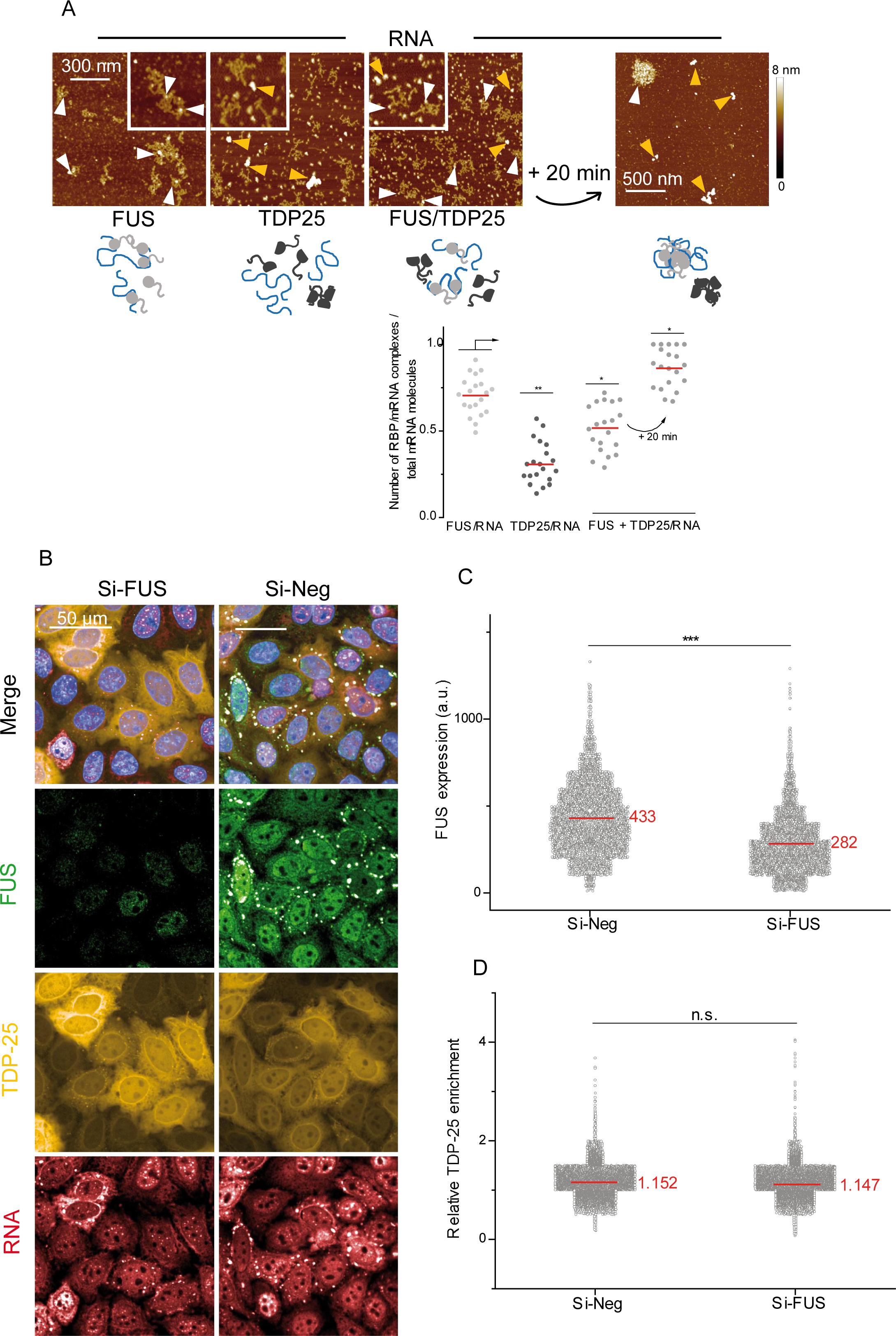
FUS does not avoid the TDP-25 cytoplasmic aggregation. (A) Top: Atomic force microscopy images (and zooms in) of FUS/RNA, TDP25/RNA and a mixture of the two RBPs with 2Luc RNA. Proteins (500 nM) were incubated with 2Luc RNA (protein/nucleotide ratio of 1/100) for 10 minutes (or additional 20 minutes for the protein mix) before sample deposition and fixation. Orange arrows pointed protein aggregates apparently free of RNA, white arrowheads protein/RNA complexes. Z scale 8 nm. Bottom: Scatter plot of the proportion of RBP/RNA complexes compared to total structures (complexes and free RNAs) adsorbed on mica surface and observed on AFM images. Each point corresponds to the same total surface analyzed, here 100 µm². The plot gathers the data from two independent experiments. Red lines show mean values. Significances between FUS/RNA, TDP25/RNA and (FUS+TDP25)/RNA samples were obtained using t test; *, p < 0.05; **, p < 0.01. (B) Representative images of HeLa cells overexpressing GFP tagged TDP-25 exposed to H2O2 and puromycin treatments to trigger SG assembly with (si-FUS) or without (si-Neg) decreasing endogenous FUS levels with siRNA. Scale bar: 50µm (C) Scatter plot representing FUS expression level in cells containing SG. Significance between expression level was obtained using t-test: ***, p < 0.005. (D) Scatter plot representing the relative TDP-25 enrichment in SGs versus cytoplasmic level in cells expressing FUS (si-Neg) or not (si-FUS). Significance between TDP-25 (relative enrichment was obtained using t-test; ns, not significant.

## Discussion

Protein aggregation is considered as deleterious for cells that have to implement molecular mechanisms to prevent the aggregation of proteins, notably for RBPs that are highly prone for aggregation since they harbor many long and self-adhesive LCDs. The presence of an elevated concentration of RNA in the nucleus is an efficient mean to prevent nuclear RBP aggregation [63]. Another mean could be provided by post-translational modifications such as phosphorylation, which modulates the solubility of RBPs. In the case of FUS, PrLD is enriched in serine/threonine residues which have a high potential of being phosphorylated to decreases theself-adhesive FUS intermolecular interactions as reported *in vitro* [64]. For TDP 43, the acetylation and phosphorylation of PrLD has been considered as a factor favoring its aggregation [65, 66]. But a recent study indicates that, as for FUS, the phosphorylation of PrLD can be favorable to the solubilization of TDP-43 [67]. Finally, cells can also use proteins playing the role of chaperones to prevent the aggregation of RBPs. The typical example is found in the interplay between aggregation-prone RBPs displaying proline-rich domains (PRMs) and soluble proteins harboring multiple SRC homology 3 domains [42]. In this case, the buffering efficiency relies on the relative SH3 and PRM domain concentration [68, 69].

Here we analyzed the consequence of an interplay between TDP-43 and FUS on TDP-43 solubility. TDP-43 is a highly aggregation-prone protein *in vitro* with a high occurrence in many neuronal cytoplasmic inclusions in neurodegenerative diseases. On the other hand, even if some FUS mutations also lead to its aggregation, the occurrence of FUS-positive inclusions in neurons is significantly less important. Accordingly, FUS is comparatively more soluble in agreement with its widespread use as a model to study phase separation *in vitro*. Besides their differential tendency for aggregation, TDP-43 and FUS share a common involvement in many stages of the mRNA metabolism but have not demonstrated an established overlap regarding their respective interactome. While many hnRNPs are present in the FUS interactome, TDP-43 is not considered as a major interactor [28, 29] or is absent from the FUS interactome [70, 71]. For the TDP-43 interactome, the observations are also divergent as studies reported either the presence [27] or absence of FUS [45]. We demonstrated here that these two proteins have a strong propensity to interact in a cellular context and this interaction is higher to what was observed with other nuclear or cytoplasmic RBPs (Figure 1B and C). It is therefore possible that this interaction offers a possibility for cells to maintain a sustainable level of solubility for TDP-43. However, as the MT bench assay works in cellular context in the presence of native RNA, it is then not possible to determine whether the interaction is direct between FUS and TDP-43 or indirect, i.e. involving another compound such as RNA or a common protein partner. To address this point, the experiments were carried out *in vitro* to confirm that TDP-43 and FUS have an inherent ability to interact with each other (Figure 9 top). Interestingly the interaction between TDP-43 and FUS is reinforced by the presence of RNA. A direct consequence of this interaction is that TDP-43 preserves its functional binding to RNA inside FUS:RNA assemblies. Indeed, even if only a small fraction of TDP-43 is incorporated in FUS assemblies, it seems to be enough to reorganize the distribution of TDP-43 on RNA (Figure 5B and E). TDP-43 binds specifically to GU sequences with a high cooperativity while FUS does not show a high sequence specificity. In a context of fierce competition between nuclear RBPs for the management of RNAs, the crosstalk between these two RBPs could favor the access of TDP-43 to its specific sequences (Figure 9 bottom left). Indeed, TDP-43 could be recruited in the FUS-rich phases along mRNA and thus, by 2D diffusion along the mRNA, gradually reach its specific sites. Note that the RNA remodeling activity associated to FUS RGGs promote the destabilization of structured RNA [34, 35], which may further increase the accessibility of TDP-43 to its specific sites. In agreement with the proposed model, FUS binds nascent RNAs, most likely in association with other FET family members (EWSR1 and TAF15) which displayed similar binding profiles on RNA [72]. The recruitment of TDP-43 to this FUS-rich phase would allow TDP-43 to gain access to its specific sites on nascent mRNAs. Interestingly, while FUS intron 7 RNA is known to be a target of FUS itself to orchestrate its self-regulation, a conserved TDP-43 binding site has also been identified in the same intron causing a reduction of FUS intron retention upon TDP-43 knockdown [73]. This result is in line with a putative interplay between TDP-43 and FUS on introns during transcription.

**Figure 9:**
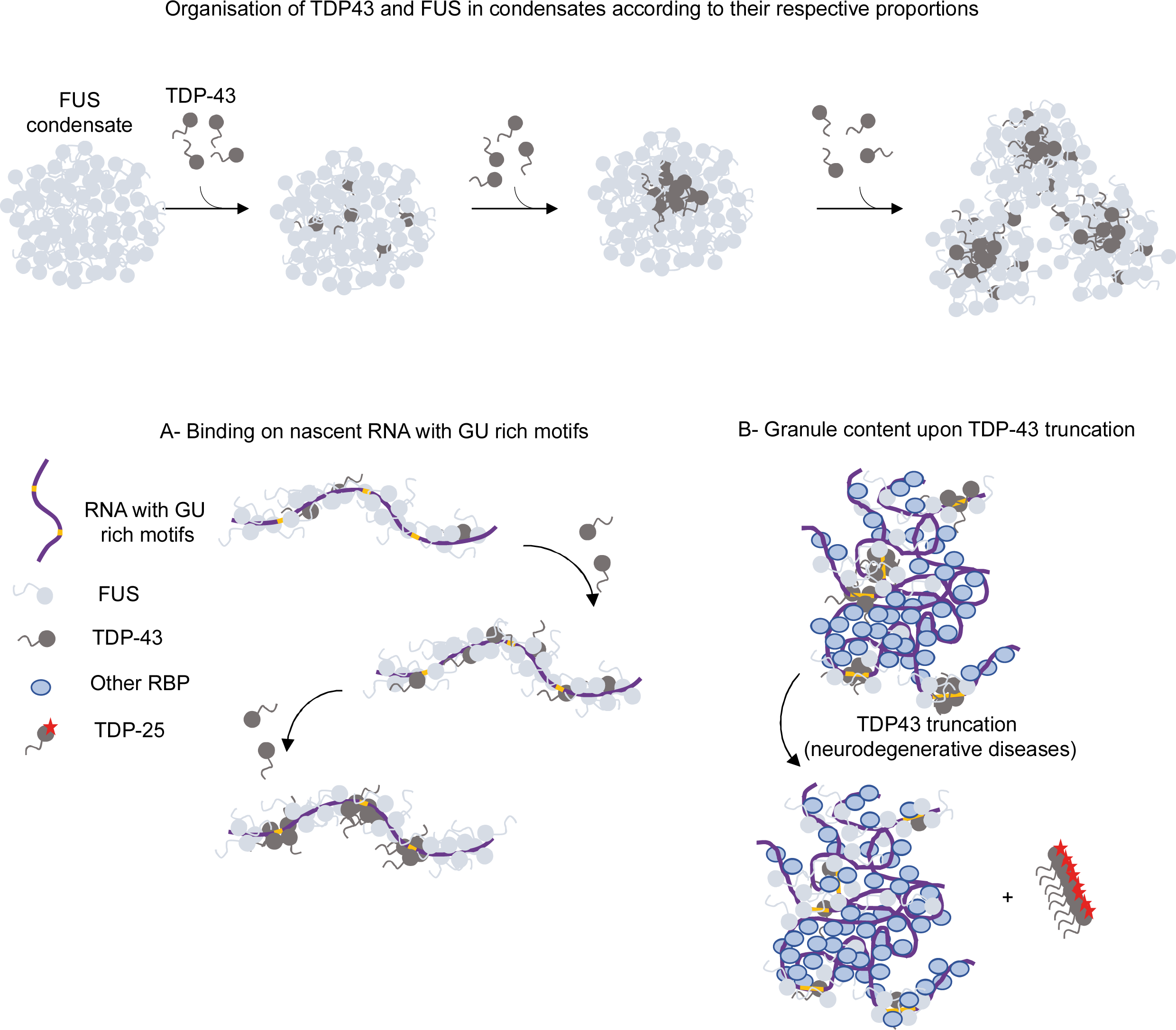
Sub-compartimentalization of TDP-43 in FUS rich-phases. Top: *in vitro*, without RNA, FUS assembles into condensate in which few TDP-43 molecules could be solubilized representing a typical example of sub-compartmentalization. Additional TDP-43 units could integrate the FUS condensate until the solubilization limit is reached. Beyond, additional TDP-43 form sub-compartments and then promotes the coalescence of the few condensates leading to large aggregates with a filamentous appearance. The solubilization of TDP-43 into FUS rich phases may favor the accessibility of nascent mRNA to TDP-43 and its specific binding to GU rich sequences (bottom, left). When TDP-43 loses its mRNA binding ability (ex. truncated TDP-25), the multivalent interactions between PrLD of both proteins are not sufficient to maintain the mixing with FUS. It could promote the exclusion of TDP-25 from mRNA rich granules like stress granules and the formation of TDP-25 insoluble aggregates (bottom, right).

If FUS accepts TDP-43 in its environment and participates in its solubilization, the reverse is not clear. Indeed, the miscibility of FUS in the TDP-43-rich phase is very low (Figure 3H). Even in the presence of RNA, the correlation coefficient between the signals of these proteins remains low (Figure 4E), reflecting a highly limited miscibility of FUS in TDP-43-rich phase. The difference in miscibility depends on the phase considered which is a common behavior in polymers that have partial miscibility [74]. The difficulty for FUS to mix with the TDP-43-rich phase can also be related to the structure of the RBP higher-order assemblies. Indeed, while FUS is organized in small spherical aggregates with a very strong resemblance to droplets, TDP-43 forms massive assemblies with a filamentous appearance (Figure 3A and B). By analogy with organic polymers, one can suspect a rather amorphous structure in the assemblies of FUS in which the mobility of the molecules would make it possible to accept a certain proportion of TDP-43 in connection also with the porous structure of the hydrogels formed by FUS [75] in which some RBPs like STAU1 or SMN can diffuse [53]. TDP-43 assemblies would be closer to semi crystalline polymers whose structure of the dense crystalline phase prohibits any mixture with another polymer. Moreover, the mobility of molecules measured by FRAP in FUS aggregates is much higher than that measured in assemblies of TDP-43 [76].

To elaborate about the putative perspectives of the results presented here in neurodegeneration, we consider that TDP-43 cytoplasmic inclusions are hallmarks of almost all cases of ALS and half of FTD case. In contrast, FUS inclusions are less frequent in sporadic ALS and uncommon in FTD. Considering the interaction between TDP-43 and FUS and the fact that TDP-43 could be solubilized to a certain extent in the FUS-rich phases (Figure 5B and 6), one may wonder whether the presence of FUS could prevent the irreversible aggregation of TDP-43 in neurons of ALS/FTLD patients. Analysis of the composition of the cytoplasmic aggregates initially produced contradictory results, some suggesting co-aggregation of TDP-43 and FUS [77–80], others indicating distinct aggregation [30, 31, 81–84]. Here, we evidenced that TDP-43 and FUS colocalize in cytoplasmic SGs and that the overexpression of FUS favors the enrichment of TDP-43 in SGs compare to other proteins (Figure 6). In addition, using a defective cooperative-RNA binding TDP-43, the enrichment of TDP-43 as well as FUS in SGs is decreased compared to TDP-43 WT overexpression. However, analysis of the structure of pathologic cytoplasmic aggregates in patients with ALS suggests that these two proteins undergo independent aggregation processes [85]. Interestingly, the cytoplasmic localization of FUS, independently of its aggregation, is a generic hallmark of ALS case with TDP-43 cytoplasmic aggregation [86]. Thus, while these two RBPs are mainly nuclear and involved in the same processes, they could both be found in the cytoplasm, but one in a soluble form, FUS, while the other, TDP-43, upon pathological conditions, is aggregated. The aggregation of TDP-43 could therefore be linked to the disruption of its interaction with FUS on their common mRNA targets, either due to mutations or truncations (Figure 9 bottom right). Here we demonstrate that when TDP-43 loses its RNA-binding capacity as in the case of TDP-25 truncation, it is no longer in the environment of FUS, which promotes TDP-25 aggregation. TDP-25 is widely represented in cytoplasmic aggregates in the brain of both FTLD and ALS patients [87–89] and the loss of functional TDP-43 and FUS interplay could participate in the cytoplasmic aggregation of TDP-25.

From the data at different scales, *in vitro* and in cells presented herein, we have demonstrated that the interplay between TDP-43 and FUS allows TDP-43 to be recruited in FUS-rich phases. TDP-43 recruitment helps to preserve TDP-43 interaction with RNA and enables the formation of TDP-43-rich sub-compartments. This relationship opens news perspectives to better understand the mechanisms by which TDP-43 forms aggregates in a pathological context. Indeed, while most cytoplasmic inclusions are TDP-43-positive in ALS and FTLD, FUS mutations induce the formation of FUS-rich inclusions but not that of TDP-43, which would have to be the case if the binding of TDP-43 to RNA would rely solely on FUS. Indeed, one should bear in minds that there are other abundant FET proteins, TAF15 and EWSR1, which may also recruit TDP-43 in their compartments to compensate for the loss of FUS, which deserves to be explored.

## Acknowledgements

We gratefully acknowledge the Genopole Evry, the University of Evry and INSERM for constant support of the laboratory. This study was also supported by the Doctoral School of the University Paris Saclay (Grant for CD). We thank also Pr Dieter Edbauer from the Deutsches Zentrum für Neurodegenerative Erkrankungen in Munich-Germany for providing their TDP-25 plasmid.

## Author contributions

C.D., S.T., D.P. and L.H. conceived and planned the experiments. C.D. carried out the experiments. V.J., L.C. and B.D. contributed to sample preparation. C.D., S.T., D.P. and L.H discussed the results and commented on the manuscript. D.P. contributed to the final version of the manuscript. DP and L.H. conceived the project. L.H. wrote the manuscript.

## Data availability

All the data presented in this study are available on request from the corresponding author.

## Competing interests

The authors declare no competing interests

Correspondence and requests for materials should be addressed to L.H.

## References

[1] Arai T, Hasegawa M, Akiyama H, Ikeda K, Nonaka T, Mori H, et al. TDP-43 is a component of ubiquitin-positive tau-negative inclusions in frontotemporal lobar degeneration and amyotrophic lateral sclerosis. Biochemical and biophysical research communications 2006;351:602–11.

[2] Neumann M, Sampathu DM, Kwong LK, Truax AC, Micsenyi MC, Chou TT, et al. Ubiquitinated TDP-43 in frontotemporal lobar degeneration and amyotrophic lateral sclerosis. Science 2006;314:130–3.

[3] Kwiatkowski TJ, Jr., Bosco DA, Leclerc AL, Tamrazian E, Vanderburg CR, Russ C, et al. Mutations in the FUS/TLS gene on chromosome 16 cause familial amyotrophic lateral sclerosis. Science 2009;323:1205–8.

[4] Vance C, Rogelj B, Hortobagyi T, De Vos KJ, Nishimura AL, Sreedharan J, et al. Mutations in FUS, an RNA processing protein, cause familial amyotrophic lateral sclerosis type 6. Science 2009;323:1208–11.

[5] Klim JR, Pintacuda G, Nash LA, Guerra San Juan I, Eggan K. Connecting TDP-43 Pathology with Neuropathy. Trends in neurosciences 2021;44:424–40.

[6] Ratti A, Buratti E. Physiological functions and pathobiology of TDP-43 and FUS/TLS proteins. Journal of neurochemistry 2016;138 Suppl 1:95–111.

[7] Deng H, Gao K, Jankovic J. The role of FUS gene variants in neurodegenerative diseases. Nature reviews Neurology 2014;10:337–48.

[8] Maharana S, Wang J, Papadopoulos DK, Richter D, Pozniakovsky A, Poser I, et al. RNA buffers the phase separation behavior of prion-like RNA binding proteins. Science 2018;360:918–21.

[9] Mann JR, Gleixner AM, Mauna JC, Gomes E, DeChellis-Marks MR, Needham PG, et al. RNA Binding Antagonizes Neurotoxic Phase Transitions of TDP-43. Neuron 2019;102:321–38 e8.

[10] Bounedjah O, Desforges B, Wu TD, Pioche-Durieu C, Marco S, Hamon L, et al. Free mRNA in excess upon polysome dissociation is a scaffold for protein multimerization to form stress granules. Nucleic acids research 2014;42:8678–91.

[11] Zacco E, Grana-Montes R, Martin SR, de Groot NS, Alfano C, Tartaglia GG, et al. RNA as a key factor in driving or preventing self-assembly of the TAR DNA-binding protein 43. Journal of molecular biology 2019;431:1671–88.

[12] Li YR, King OD, Shorter J, Gitler AD. Stress granules as crucibles of ALS pathogenesis. The Journal of cell biology 2013;201:361–72.

[13] Zhang P, Fan B, Yang P, Temirov J, Messing J, Kim HJ, et al. Chronic optogenetic induction of stress granules is cytotoxic and reveals the evolution of ALS-FTD pathology. eLife 2019;8.

[14] Chen HJ, Topp SD, Hui HS, Zacco E, Katarya M, McLoughlin C, et al. RRM adjacent TARDBP mutations disrupt RNA binding and enhance TDP-43 proteinopathy. Brain : a journal of neurology 2019;142:3753–70.

[15] Daigle JG, Lanson NA, Jr., Smith RB, Casci I, Maltare A, Monaghan J, et al. RNA-binding ability of FUS regulates neurodegeneration, cytoplasmic mislocalization and incorporation into stress granules associated with FUS carrying ALS-linked mutations. Human molecular genetics 2013;22:1193–205.

[16] Ling SC, Polymenidou M, Cleveland DW. Converging mechanisms in ALS and FTD: disrupted RNA and protein homeostasis. Neuron 2013;79:416–38.

[17] Tollervey JR, Curk T, Rogelj B, Briese M, Cereda M, Kayikci M, et al. Characterizing the RNA targets and position-dependent splicing regulation by TDP-43. Nature neuroscience 2011;14:452–8.

[18] Lagier-Tourenne C, Polymenidou M, Hutt KR, Vu AQ, Baughn M, Huelga SC, et al. Divergent roles of ALS-linked proteins FUS/TLS and TDP-43 intersect in processing long pre-mRNAs. Nature neuroscience 2012;15:1488–97.

[19] Rogelj B, Easton LE, Bogu GK, Stanton LW, Rot G, Curk T, et al. Widespread binding of FUS along nascent RNA regulates alternative splicing in the brain. Scientific reports 2012;2:603.

[20] Honda D, Ishigaki S, Iguchi Y, Fujioka Y, Udagawa T, Masuda A, et al. The ALS/FTLD-related RNA-binding proteins TDP-43 and FUS have common downstream RNA targets in cortical neurons. FEBS open bio 2013;4:1–10.

[21] Kim SH, Shanware NP, Bowler MJ, Tibbetts RS. Amyotrophic lateral sclerosis-associated proteins TDP-43 and FUS/TLS function in a common biochemical complex to co-regulate HDAC6 mRNA. The Journal of biological chemistry 2010;285:34097–105.

[22] Kabashi E, Bercier V, Lissouba A, Liao M, Brustein E, Rouleau GA, et al. FUS and TARDBP but not SOD1 interact in genetic models of amyotrophic lateral sclerosis. PLoS genetics 2011;7:e1002214.

[23] Lanson NA, Jr., Maltare A, King H, Smith R, Kim JH, Taylor JP, et al. A Drosophila model of FUS-related neurodegeneration reveals genetic interaction between FUS and TDP-43. Human molecular genetics 2011;20:2510–23.

[24] Wang JW, Brent JR, Tomlinson A, Shneider NA, McCabe BD. The ALS-associated proteins FUS and TDP-43 function together to affect Drosophila locomotion and life span. The Journal of clinical investigation 2011;121:4118–26.

[25] Avendano-Vazquez SE, Dhir A, Bembich S, Buratti E, Proudfoot N, Baralle FE. Autoregulation of TDP-43 mRNA levels involves interplay between transcription, splicing, and alternative polyA site selection. Genes & development 2012;26:1679–84.

[26] Masuda A, Takeda J, Okuno T, Okamoto T, Ohkawara B, Ito M, et al. Position-specific binding of FUS to nascent RNA regulates mRNA length. Genes & development 2015;29:1045–57.

[27] Ling SC, Albuquerque CP, Han JS, Lagier-Tourenne C, Tokunaga S, Zhou H, et al. ALS-associated mutations in TDP-43 increase its stability and promote TDP-43 complexes with FUS/TLS. Proceedings of the National Academy of Sciences of the United States of America 2010;107:13318–23.

[28] Reber S, Jutzi D, Lindsay H, Devoy A, Mechtersheimer J, Levone BR, et al. The phase separation-dependent FUS interactome reveals nuclear and cytoplasmic function of liquid-liquid phase separation. Nucleic acids research 2021;49:7713–31.

[29] Sun S, Ling SC, Qiu J, Albuquerque CP, Zhou Y, Tokunaga S, et al. ALS-causative mutations in FUS/TLS confer gain and loss of function by altered association with SMN and U1-snRNP. Nature communications 2015;6:6171.

[30] Sun Z, Diaz Z, Fang X, Hart MP, Chesi A, Shorter J, et al. Molecular determinants and genetic modifiers of aggregation and toxicity for the ALS disease protein FUS/TLS. PLoS biology 2011;9:e1000614.

[31] Urwin H, Josephs KA, Rohrer JD, Mackenzie IR, Neumann M, Authier A, et al. FUS pathology defines the majority of tau- and TDP-43-negative frontotemporal lobar degeneration. Acta neuropathologica 2010;120:33–41.

[32] Schwartz JC, Wang X, Podell ER, Cech TR. RNA seeds higher-order assembly of FUS protein. Cell reports 2013;5:918–25.

[33] Ozdilek BA, Thompson VF, Ahmed NS, White CI, Batey RT, Schwartz JC. Intrinsically disordered RGG/RG domains mediate degenerate specificity in RNA binding. Nucleic acids research 2017;45:7984–96.

[34] Loughlin FE, Lukavsky PJ, Kazeeva T, Reber S, Hock EM, Colombo M, et al. The Solution Structure of FUS Bound to RNA Reveals a Bipartite Mode of RNA Recognition with Both Sequence and Shape Specificity. Molecular cell 2019;73:490–504 e6.

[35] Loughlin FE, Wilce JA. TDP-43 and FUS-structural insights into RNA recognition and self-association. Current opinion in structural biology 2019;59:134–42.

[36] Polymenidou M, Lagier-Tourenne C, Hutt KR, Huelga SC, Moran J, Liang TY, et al. Long pre-mRNA depletion and RNA missplicing contribute to neuronal vulnerability from loss of TDP-43. Nature neuroscience 2011;14:459–68.

[37] Lukavsky PJ, Daujotyte D, Tollervey JR, Ule J, Stuani C, Buratti E, et al. Molecular basis of UG-rich RNA recognition by the human splicing factor TDP-43. Nature structural & molecular biology 2013;20:1443–9.

[38] Rengifo-Gonzalez JC, El Hage K, Clement MJ, Steiner E, Joshi V, Craveur P, et al. The cooperative binding of TDP-43 to GU-rich RNA repeats antagonizes TDP-43 aggregation. eLife 2021;10.

[39] Maucuer A, Desforges B, Joshi V, Boca M, Kretov D, Hamon L, et al. Microtubules as platforms for probing liquid-liquid phase separation in cells: application to RNA-binding proteins. Journal of cell science 2018.

[40] Abrakhi S, Kretov DA, Desforges B, Dobra I, Bouhss A, Pastre D, et al. Nanoscale Analysis Reveals the Maturation of Neurodegeneration-Associated Protein Aggregates: Grown in mRNA Granules then Released by Stress Granule Proteins. ACS nano 2017;11:7189–200.

[41] Boca M, Kretov DA, Desforges B, Mephon-Gaspard A, Curmi PA, Pastre D. Probing protein interactions in living mammalian cells on a microtubule bench. Scientific reports 2015;5:17304.

[42] Pankivskyi S, Pastre D, Steiner E, Joshi V, Rynditch A, Hamon L. ITSN1 regulates SAM68 solubility through SH3 domain interactions with SAM68 proline-rich motifs. Cellular and molecular life sciences : CMLS 2020.

[43] Singatulina AS, Hamon L, Sukhanova MV, Desforges B, Joshi V, Bouhss A, et al. PARP-1 Activation Directs FUS to DNA Damage Sites to Form PARG-Reversible Compartments Enriched in Damaged DNA. Cell reports 2019;27:1809–21 e5.

[44] Samsonova A, El Hage K, Desforges B, Joshi V, Clement MJ, Lambert G, et al. Lin28, a major translation reprogramming factor, gains access to YB-1-packaged mRNA through its cold-shock domain. Communications biology 2021;4:359.

[45] Freibaum BD, Chitta RK, High AA, Taylor JP. Global analysis of TDP-43 interacting proteins reveals strong association with RNA splicing and translation machinery. Journal of proteome research 2010;9:1104–20.

[46] McCarty J, Delaney KT, Danielsen SPO, Fredrickson GH, Shea JE. Complete Phase Diagram for Liquid-Liquid Phase Separation of Intrinsically Disordered Proteins. J Phys Chem Lett 2019;10:1644–52.

[47] Meyer NH, Tripsianes K, Vincendeau M, Madl T, Kateb F, Brack-Werner R, et al. Structural basis for homodimerization of the Src-associated during mitosis, 68-kDa protein (Sam68) Qua1 domain. The Journal of biological chemistry 2010;285:28893–901.

[48] Feracci M, Foot JN, Grellscheid SN, Danilenko M, Stehle R, Gonchar O, et al. Structural basis of RNA recognition and dimerization by the STAR proteins T-STAR and Sam68. Nature communications 2016;7:10355.

[49] Chen T, Boisvert FM, Bazett-Jones DP, Richard S. A role for the GSG domain in localizing Sam68 to novel nuclear structures in cancer cell lines. Molecular biology of the cell 1999;10:3015–33.

[50] Scheiba RM, de Opakua AI, Diaz-Quintana A, Cruz-Gallardo I, Martinez-Cruz LA, Martinez-Chantar ML, et al. The C-terminal RNA binding motif of HuR is a multi-functional domain leading to HuR oligomerization and binding to U-rich RNA targets. RNA biology 2014;11:1250–61.

[51] Wang J, Choi JM, Holehouse AS, Lee HO, Zhang X, Jahnel M, et al. A Molecular Grammar Governing the Driving Forces for Phase Separation of Prion-like RNA Binding Proteins. Cell 2018;174:688–99 e16.

[52] Murthy AC, Dignon GL, Kan Y, Zerze GH, Parekh SH, Mittal J, et al. Molecular interactions underlying liquid-liquid phase separation of the FUS low-complexity domain. Nature structural & molecular biology 2019;26:637–48.

[53] Murakami T, Qamar S, Lin JQ, Schierle GS, Rees E, Miyashita A, et al. ALS/FTD Mutation-Induced Phase Transition of FUS Liquid Droplets and Reversible Hydrogels into Irreversible Hydrogels Impairs RNP Granule Function. Neuron 2015;88:678–90.

[54] Murray DT, Kato M, Lin Y, Thurber KR, Hung I, McKnight SL, et al. Structure of FUS Protein Fibrils and Its Relevance to Self-Assembly and Phase Separation of Low-Complexity Domains. Cell 2017;171:615–27 e16.

[55] Qamar S, Wang G, Randle SJ, Ruggeri FS, Varela JA, Lin JQ, et al. FUS Phase Separation Is Modulated by a Molecular Chaperone and Methylation of Arginine Cation-pi Interactions. Cell 2018;173:720–34 e15.

[56] Murray DT, Tycko R. Side Chain Hydrogen-Bonding Interactions within Amyloid-like Fibrils Formed by the Low-Complexity Domain of FUS: Evidence from Solid State Nuclear Magnetic Resonance Spectroscopy. Biochemistry 2020;59:364–78.

[57] Monahan Z, Ryan VH, Janke AM, Burke KA, Rhoads SN, Zerze GH, et al. Phosphorylation of the FUS low-complexity domain disrupts phase separation, aggregation, and toxicity. The EMBO journal 2017;36:2951–67.

[58] Ayala YM, Zago P, D’Ambrogio A, Xu YF, Petrucelli L, Buratti E, et al. Structural determinants of the cellular localization and shuttling of TDP-43. Journal of cell science 2008;121:3778-85.

[59] Dormann D, Rodde R, Edbauer D, Bentmann E, Fischer I, Hruscha A, et al. ALS-associated fused in sarcoma (FUS) mutations disrupt Transportin-mediated nuclear import. The EMBO journal 2010;29:2841–57.

[60] Igaz LM, Kwong LK, Xu Y, Truax AC, Uryu K, Neumann M, et al. Enrichment of C-terminal fragments in TAR DNA-binding protein-43 cytoplasmic inclusions in brain but not in spinal cord of frontotemporal lobar degeneration and amyotrophic lateral sclerosis. The American journal of pathology 2008;173:182–94.

[61] Afroz T, Hock EM, Ernst P, Foglieni C, Jambeau M, Gilhespy LAB, et al. Functional and dynamic polymerization of the ALS-linked protein TDP-43 antagonizes its pathologic aggregation. Nature communications 2017;8:45.

[62] Zhang YJ, Xu YF, Cook C, Gendron TF, Roettges P, Link CD, et al. Aberrant cleavage of TDP-43 enhances aggregation and cellular toxicity. Proceedings of the National Academy of Sciences of the United States of America 2009;106:7607–12.

[63] Grese ZR, Bastos AC, Mamede LD, French RL, Miller TM, Ayala YM. Specific RNA interactions promote TDP-43 multivalent phase separation and maintain liquid properties. EMBO reports 2021:e53632.

[64] Rhoads SN, Monahan ZT, Yee DS, Shewmaker FP. The Role of Post-Translational Modifications on Prion-Like Aggregation and Liquid-Phase Separation of FUS. International journal of molecular sciences 2018;19.

[65] Buratti E. TDP-43 post-translational modifications in health and disease. Expert opinion on therapeutic targets 2018;22:279–93.

[66] Keating SS, Bademosi AT, San Gil R, Walker AK. Aggregation-prone TDP-43 sequesters and drives pathological transitions of free nuclear TDP-43. Cellular and molecular life sciences : CMLS 2023;80:95.

[67] Gruijs da Silva LA, Simonetti F, Hutten S, Riemenschneider H, Sternburg EL, Pietrek LM, et al. Disease-linked TDP-43 hyperphosphorylation suppresses TDP-43 condensation and aggregation. The EMBO journal 2022;41:e108443.

[68] Li P, Banjade S, Cheng HC, Kim S, Chen B, Guo L, et al. Phase transitions in the assembly of multivalent signalling proteins. Nature 2012;483:336–40.

[69] Harmon TS, Holehouse AS, Rosen MK, Pappu RV. Intrinsically disordered linkers determine the interplay between phase separation and gelation in multivalent proteins. eLife 2017;6.

[70] Chi B, O’Connell JD, Yamazaki T, Gangopadhyay J, Gygi SP, Reed R. Interactome analyses revealed that the U1 snRNP machinery overlaps extensively with the RNAP II machinery and contains multiple ALS/SMA-causative proteins. Scientific reports 2018;8:8755.

[71] Kawaguchi T, Rollins MG, Moinpour M, Morera AA, Ebmeier CC, Old WM, et al. Changes to the TDP-43 and FUS Interactomes Induced by DNA Damage. Journal of proteome research 2020;19:360–70.

[72] Hoell JI, Larsson E, Runge S, Nusbaum JD, Duggimpudi S, Farazi TA, et al. RNA targets of wild-type and mutant FET family proteins. Nature structural & molecular biology 2011;18:1428–31.

[73] Humphrey J, Birsa N, Milioto C, McLaughlin M, Ule AM, Robaldo D, et al. FUS ALS-causative mutations impair FUS autoregulation and splicing factor networks through intron retention. Nucleic acids research 2020;48:6889–905.

[74] Thomas S, Grohens Y, Jyotishkumar P. Characterization of polymer blends: miscibility, morphology and interfaces: wiley-vch; 2014.

[75] Kato M, Han TW, Xie S, Shi K, Du X, Wu LC, et al. Cell-free formation of RNA granules: low complexity sequence domains form dynamic fibers within hydrogels. Cell 2012;149:753–67.

[76] Farrawell NE, Lambert-Smith IA, Warraich ST, Blair IP, Saunders DN, Hatters DM, et al. Distinct partitioning of ALS associated TDP-43, FUS and SOD1 mutants into cellular inclusions. Scientific reports 2015;5:13416.

[77] Cohen NR, Hammans SR, Macpherson J, Nicoll JA. New neuropathological findings in Unverricht-Lundborg disease: neuronal intranuclear and cytoplasmic inclusions. Acta neuropathologica 2011;121:421–7.

[78] Deng HX, Zhai H, Bigio EH, Yan J, Fecto F, Ajroud K, et al. FUS-immunoreactive inclusions are a common feature in sporadic and non-SOD1 familial amyotrophic lateral sclerosis. Annals of neurology 2010;67:739–48.

[79] Shan X, Chiang PM, Price DL, Wong PC. Altered distributions of Gemini of coiled bodies and mitochondria in motor neurons of TDP-43 transgenic mice. Proceedings of the National Academy of Sciences of the United States of America 2010;107:16325–30.

[80] Kryndushkin D, Wickner RB, Shewmaker F. FUS/TLS forms cytoplasmic aggregates, inhibits cell growth and interacts with TDP-43 in a yeast model of amyotrophic lateral sclerosis. Protein & cell 2011;2:223–36.

[81] Baumer D, Hilton D, Paine SM, Turner MR, Lowe J, Talbot K, et al. Juvenile ALS with basophilic inclusions is a FUS proteinopathy with FUS mutations. Neurology 2010;75:611–8.

[82] Fujita Y, Fujita S, Takatama M, Ikeda M, Okamoto K. Numerous FUS-positive inclusions in an elderly woman with motor neuron disease. Neuropathology : official journal of the Japanese Society of Neuropathology 2011;31:170–6.

[83] Blair IP, Williams KL, Warraich ST, Durnall JC, Thoeng AD, Manavis J, et al. FUS mutations in amyotrophic lateral sclerosis: clinical, pathological, neurophysiological and genetic analysis. Journal of neurology, neurosurgery, and psychiatry 2010;81:639–45.

[84] Brady OA, Meng P, Zheng Y, Mao Y, Hu F. Regulation of TDP-43 aggregation by phosphorylation and p62/SQSTM1. Journal of neurochemistry 2011;116:248–59.

[85] Ikenaka K, Ishigaki S, Iguchi Y, Kawai K, Fujioka Y, Yokoi S, et al. Characteristic Features of FUS Inclusions in Spinal Motor Neurons of Sporadic Amyotrophic Lateral Sclerosis. Journal of neuropathology and experimental neurology 2020;79:370–7.

[86] Tyzack GE, Luisier R, Taha DM, Neeves J, Modic M, Mitchell JS, et al. Widespread FUS mislocalization is a molecular hallmark of amyotrophic lateral sclerosis. Brain : a journal of neurology 2019;142:2572-80.

[87] Tsuji H, Arai T, Kametani F, Nonaka T, Yamashita M, Suzukake M, et al. Molecular analysis and biochemical classification of TDP-43 proteinopathy. Brain : a journal of neurology 2012;135:3380-91.

[88] Neumann M, Kwong LK, Lee EB, Kremmer E, Flatley A, Xu Y, et al. Phosphorylation of S409/410 of TDP-43 is a consistent feature in all sporadic and familial forms of TDP-43 proteinopathies. Acta neuropathologica 2009;117:137–49.

[89] Arai T, Hasegawa M, Nonoka T, Kametani F, Yamashita M, Hosokawa M, et al. Phosphorylated and cleaved TDP-43 in ALS, FTLD and other neurodegenerative disorders and in cellular models of TDP-43 proteinopathy. Neuropathology : official journal of the Japanese Society of Neuropathology 2010;30:170–81.

