## Supplementary files 1-11 and supplementary tables for "RNA and FUS act in concert to prevent TDP-43 spatial segregation"

**Supplementary figure legends**

**Figure S1:**

Schematic representation of the correlation coefficient calculation procedure. In brief, images of the bait and the prey were merged into a single red-green image. A line crossing apparent microtubules was added and fluorescence intensity profiles along the line were generated for each channel. The line profiles were transformed into numerical values by obtaining a list of the fluorescence intensities along the line. The acquired lists were used to calculate the correlation coefficient between fluorescence intensities from both channels. At least three lines and more than ten cells from two independent experiments were analyzed for each bait-prey couple and used in a subsequent statistics analysis.

**Figure S2**

Example of U2OS cells co-transfected with the constructions encoding TDP-43 (left handed panel) or FUS (right handed panel) fused to RFP-MBD and the plasmid expressing different RBPs fused to GFP.

**Figure S3**

Upper panel: Representative images for proximity ligation assay (PLA) revealing the co-localization of TDP-43 and FUS in HeLa cells. Cells were fixed, incubated with anti-FUS and anti-TDP-43 antibodies and incubated with oligonucleotides probes, ligase and polymerase as described in the Materials and Methods section. Lower panel: scatter plot representing the ratio between the area of the PLA signal in the nucleus and the area of the nucleus. n: number of nuclei analyzed. Red lines show mean values. For each conditions, the percentage of cells with a PLA signal is indicating. ***, $p < 0.005$; t-test.

**Figure S4**

Schematic representation of the determination coefficient ($R^2$) procedure calculation using the MT bench assay when two RBPs are brought on the MTs.

**Figure S5**

Left handed panel: scatter plot representing the mixing between FUS and HuR, both fused to MBD according to the expression level of each construct. Each data point represents a value of determination coefficient $R^2$ calculated for one cell as described in the Materials and Methods section and supplementary figure S4. Representative images for a low (left handed panel) and a high (right handed panel) FUS/HuR expression level ratio.

Right handed panel: same as above but for the mixing between FUS and SAM68.

**Figure S6**

Top: Images of FUS assemblies revealed by immunofluorescence after incubation of FUS protein at different concentrations for 2 hours. Scale bar: 40 µm. Scatter plot representing the circularity of FUS assemblies for different concentrations. The plot shows the data from two independent experiments. n: number of assemblies analyzed. Red lines show mean values. Significances between circularity of FUS assemblies were obtained using t test; *, $p < 0.05$; **, $p < 0.01$.

Bottom: the same but for TDP-43 assemblies.

**Figure S7**

Process for the determination of the proportion of RBP/RNA complexes and their area from AFM image analyzes. Top panel: in this example, 22 RNA molecules were identified on the surface but only 2 have a maximum height above 2 nm (ratio of 0.09). Areas of each molecule portion detected after applying this threshold of 2 nm were determined using the particle analysis tool of the Nanoscope Analysis software (version 1.70) from Bruker. Middle panel: here, among the 27 molecules or complexes detected, 18 (blue arrowheads) are considered as RNA/protein complexes since their maximum height is above the threshold value of 2 nm. Areas of the complex detected were plotted. Bottom panel: when TDP43 is mixed with RNA, the proportion of complexes is lower than with FUS (0.17 and 0.67 respectively) but the areas of the complex portion with a height higher than 2 nm (blue arrowheads) increase.

**Figure S8**

RNA mobility shift assay duplicate of TDP-43 and FUS interacting together or independently with RNA. 115 ng of 2Luc RNA were mixed for 10 min with preincubated (50 min) TDP-43 (fixed at 3 µM) or FUS (concentration ranging between 0.1 to 9 µM) or a mix of both. Amounts of RNA interacting with RBP were quantified for each lane and compared for the mix (lane 8-12, grey circles) with the sum of lane 2 plus lane 3 to 7 (grey diamonds). Arrows highlight conditions were the amount of RNA interacting with RBP is higher when RBPs are mixed than considered independently

**Figure S9**

Representative images of U2OS cells co transfected with different TDP-43 constructs and FUS FL revealing diverse degrees of mixing in comparison with cell transfected with RFP and GFP-TDP-43-MBD. Scale bar: 15 µm.

**Figure S10**

Top: schematic representation of the domains of FUS truncations used in the microtubule bench assay to reveal their mixing with full length TDP-43. Bottom: scatter plot representing the mixing between TDP-43 full length and FUS truncated forms, both fused to MBD. Each data point represents a value of determination coefficient $R^2$ calculated for one cell. Red lines show mean values. Significances between determination coefficients were obtained using t test; **, $p < 0.01$.

**Figure S11**

Scatter plot representing the relative enrichment of RNA in stress granules compared to the cytoplasm in cells with a normal (Si-Neg) or low (Si-FUS) FUS expression level. Significance was obtained using t test; ns: non significant.

**Supplementary table**

1: table of plasmids used for the MT bench bait-prey method

2: table of plasmids used for the MT bench compartmentalization method.

3: table of recombinant proteins produced for *in vitro* experiments and table of plasmids for production of recombinant proteins.

4: table of primary antibodies used for immunofluorescence experiences.

**Supplementary Video**

SV1: Three dimensional reconstruction of compiled confocal images of a large assembly of TDP-43 obtained after incubation for 2h.

SV2: Three dimensional reconstruction of compiled confocal images of assembly corresponding to Figure 4C obtained after incubation of FUS (in red) and TDP-43 (in green) at a FUS/TDP-43 molar ratio of 3/1 in presence of 2Luc mRNA (protein/nucleotide = 1/10) .

SV3: Same as SV2 with a TDP-43/FUS molar ratio of 9/1 (Figure 4D)

**Calculation of the correlation coefficient from the bait/prey method (MT bench experiment)**

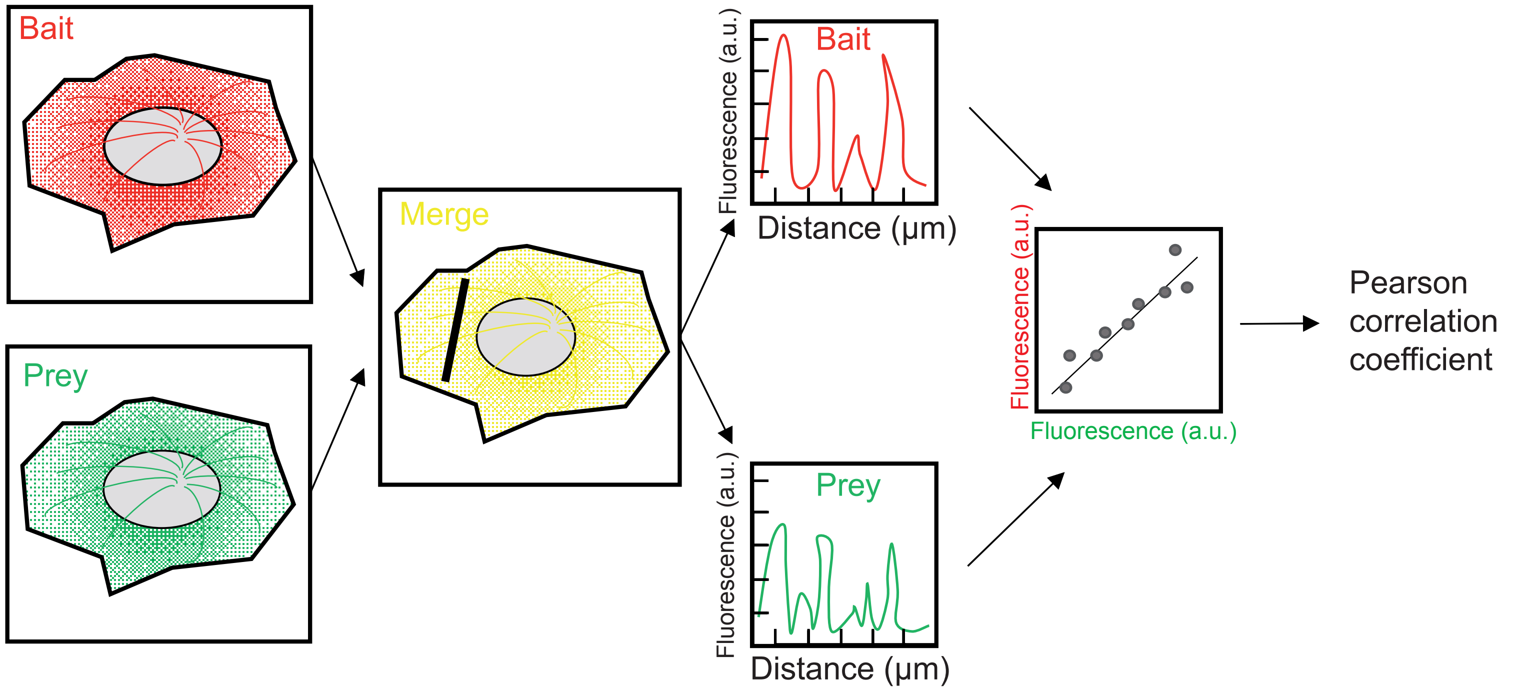


1. Aquisition of images of cells in red and green channels

2. A line crossing microtubules is drawn on merged image

3. Fluorescence intensity along the line is measured and transformed into numerical values

4. Pearson correlation coefficient is calculated for the two data sets

Supplementary figure S1

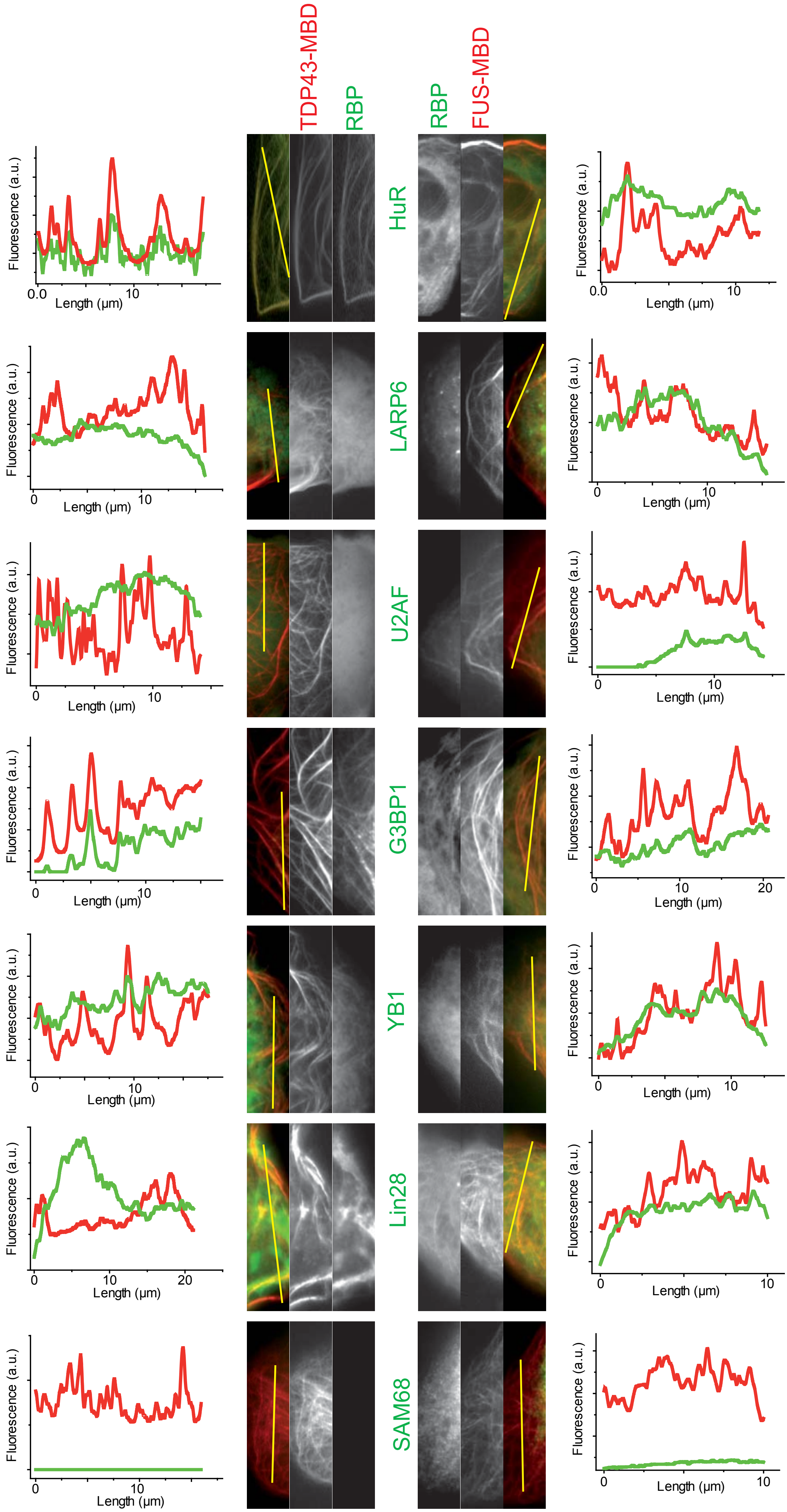


Supplementary figure S2

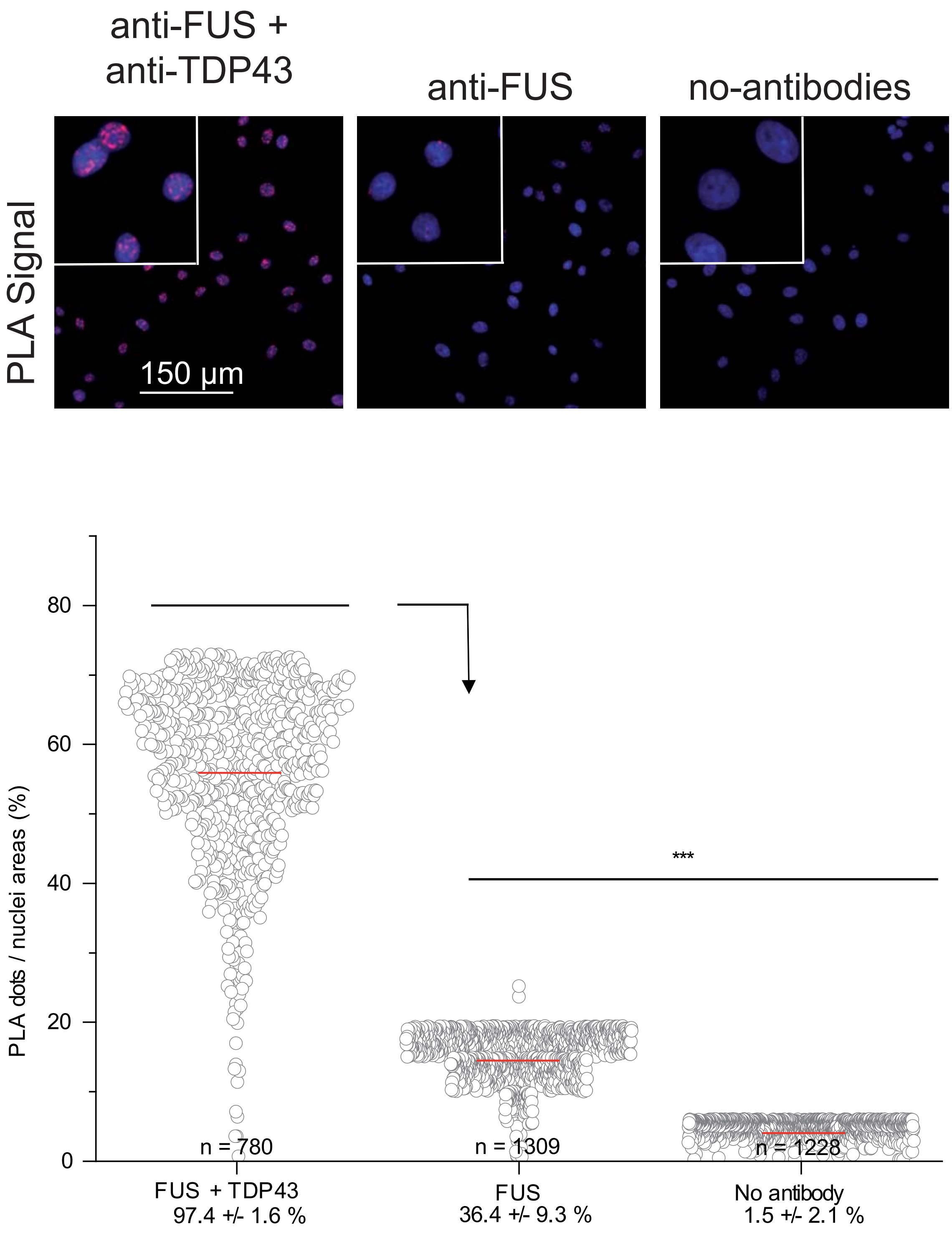


Supplementary Figure S3

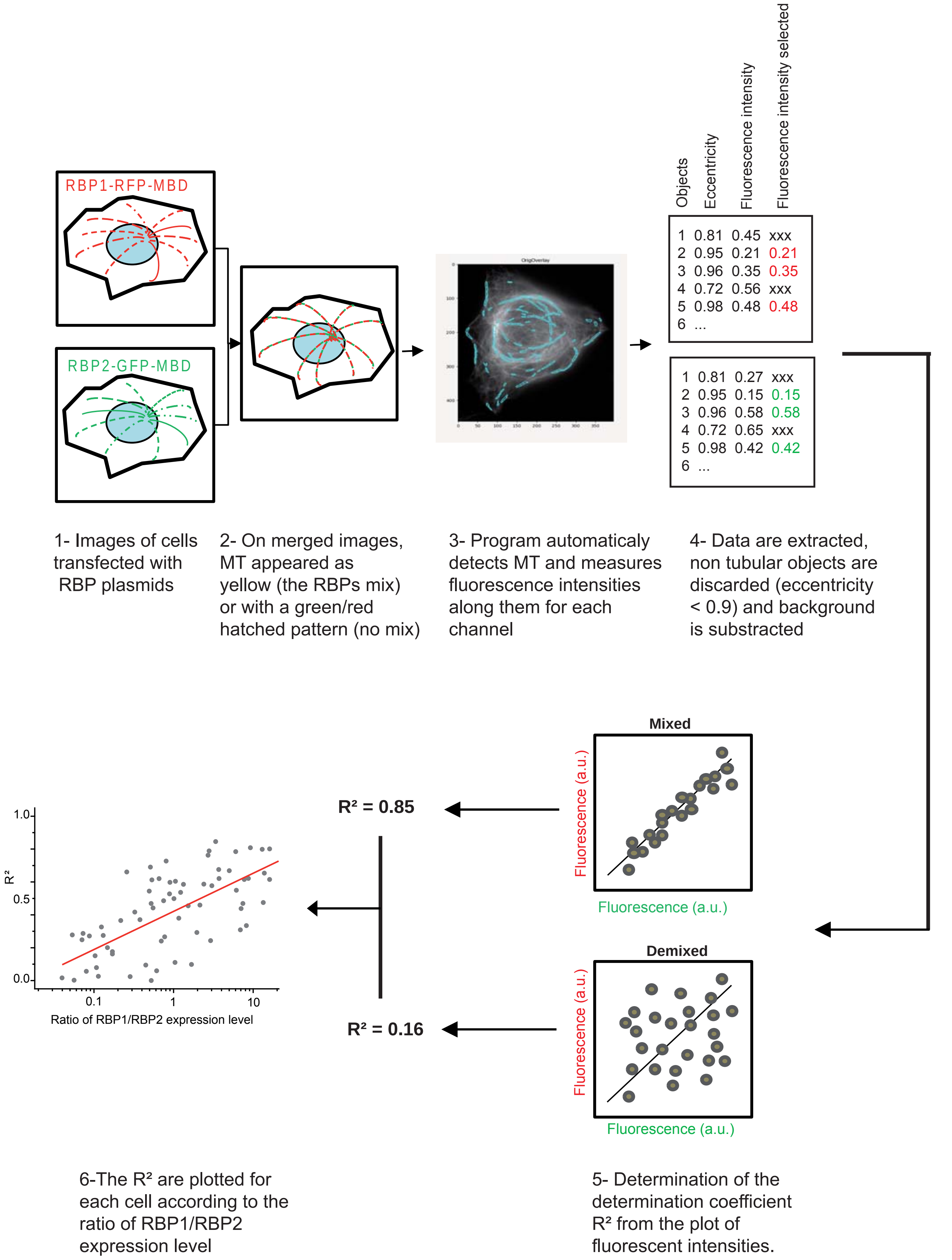


Supplementary Figure S4

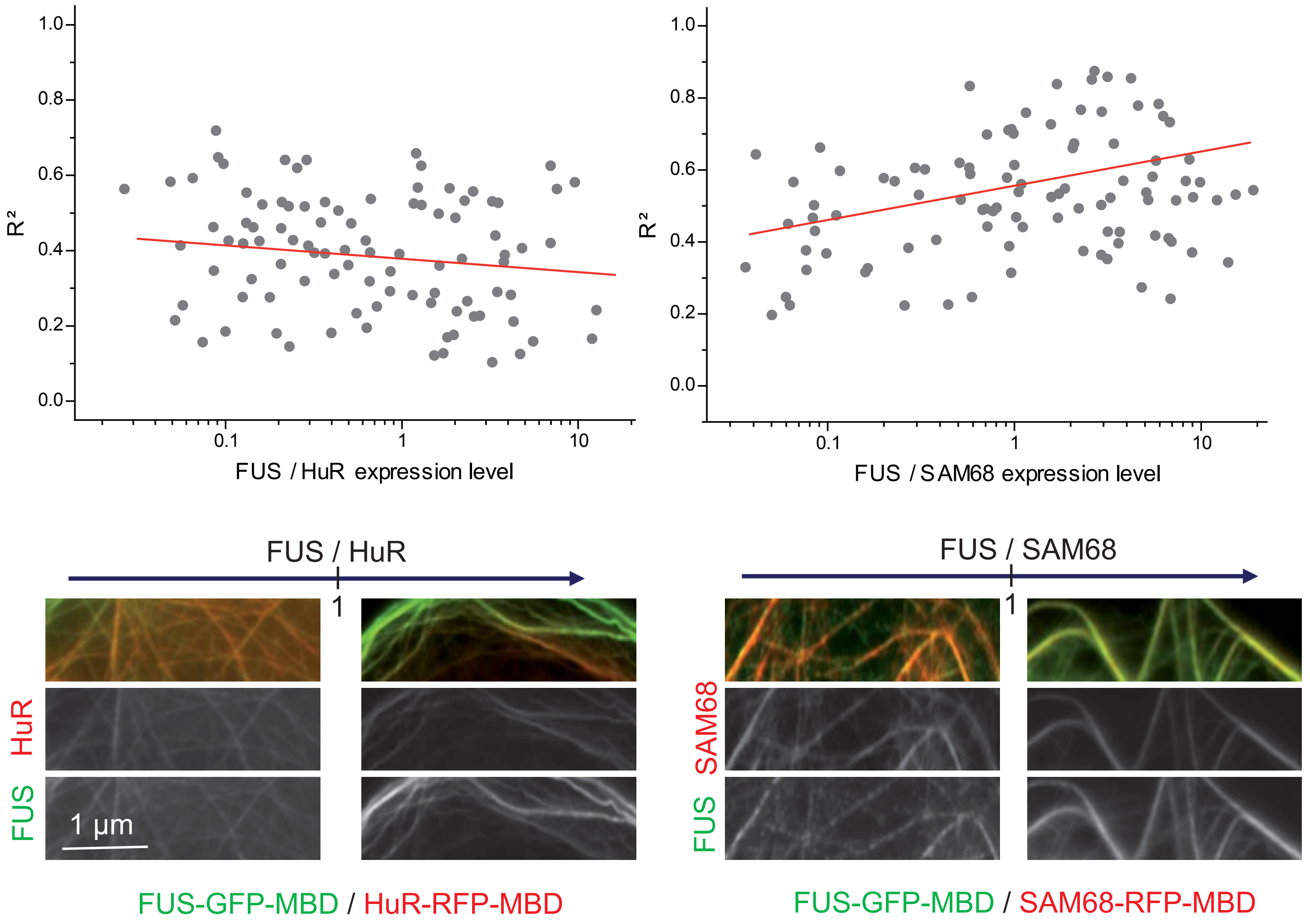


Supplementary figure S5

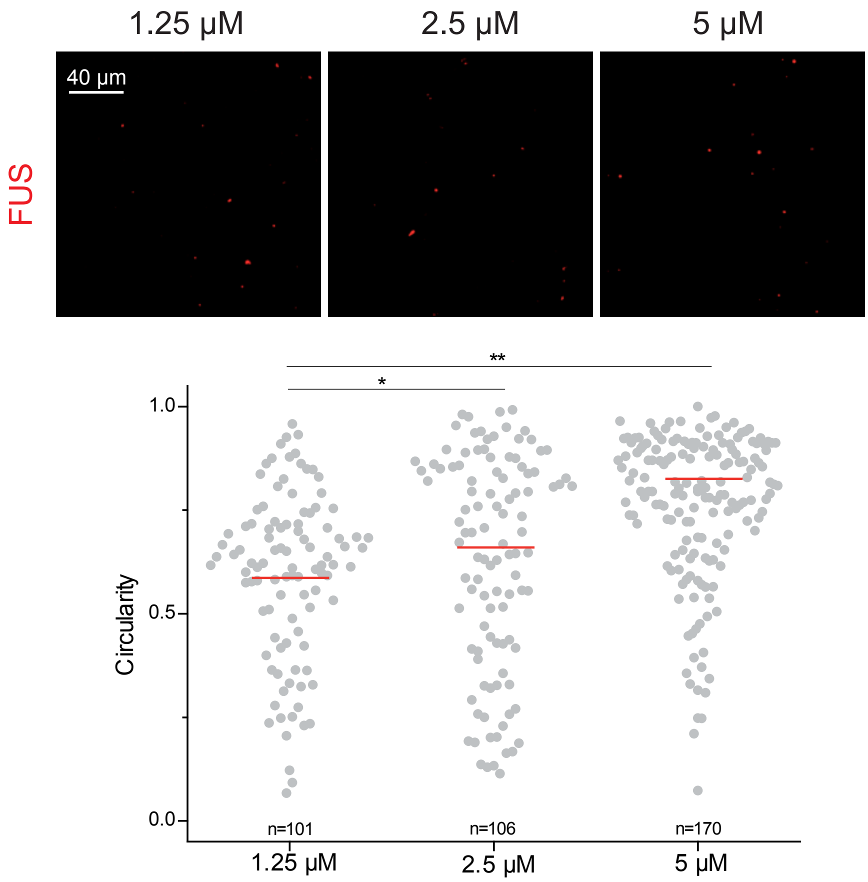


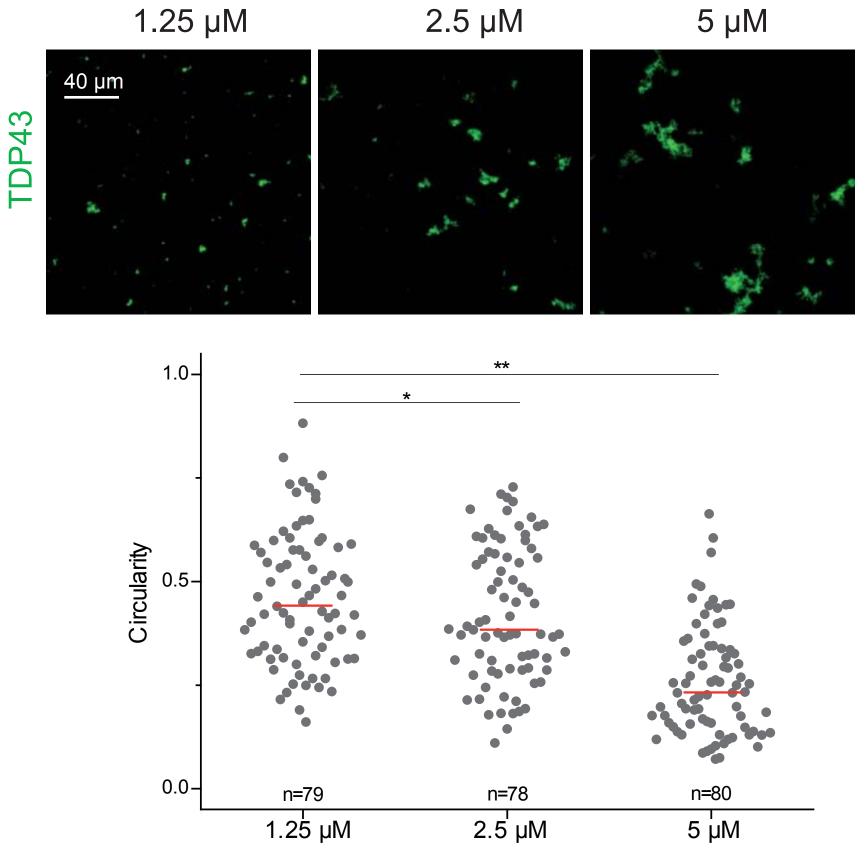


Supplementary Figure S6

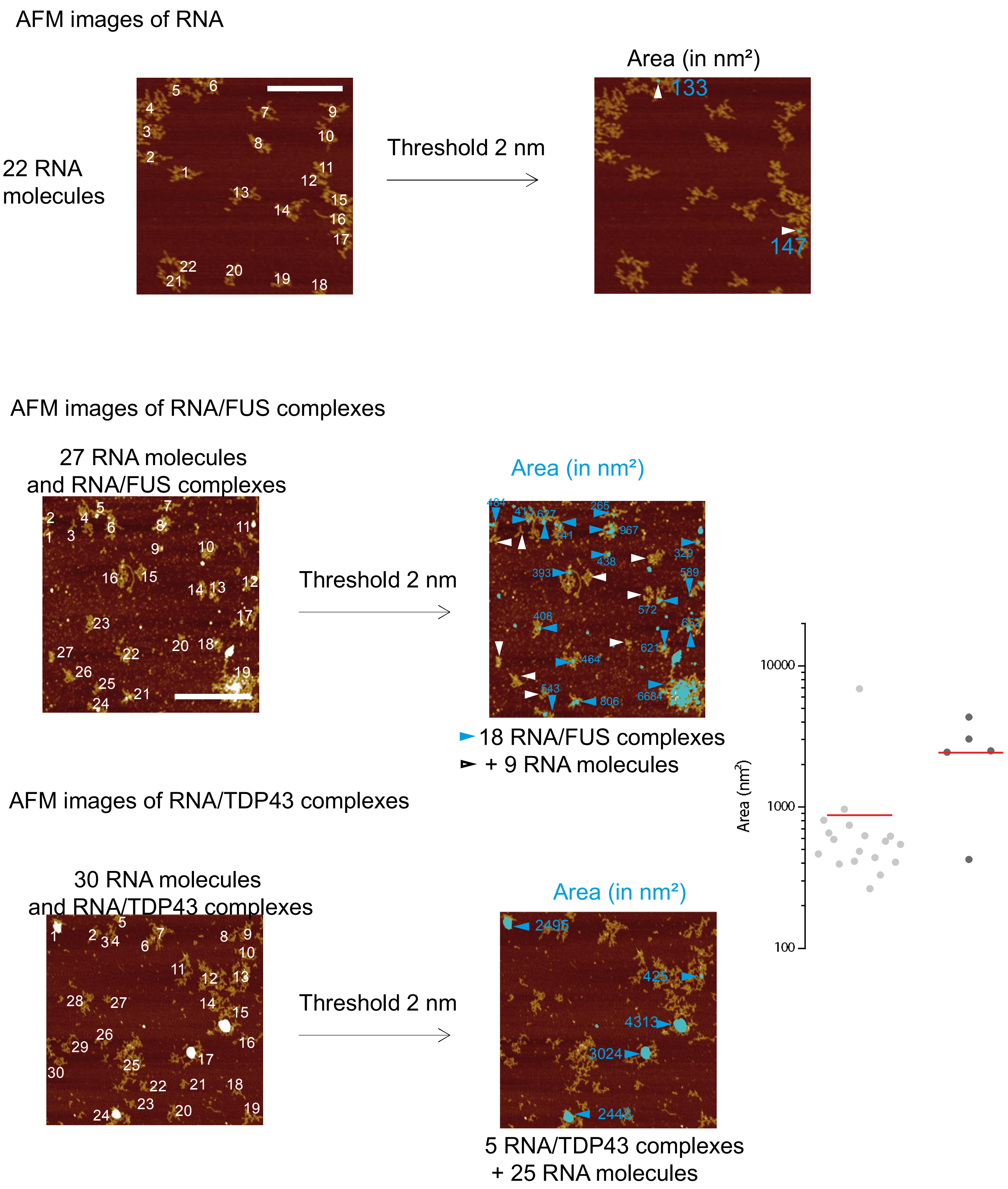


Supplementary figure S7

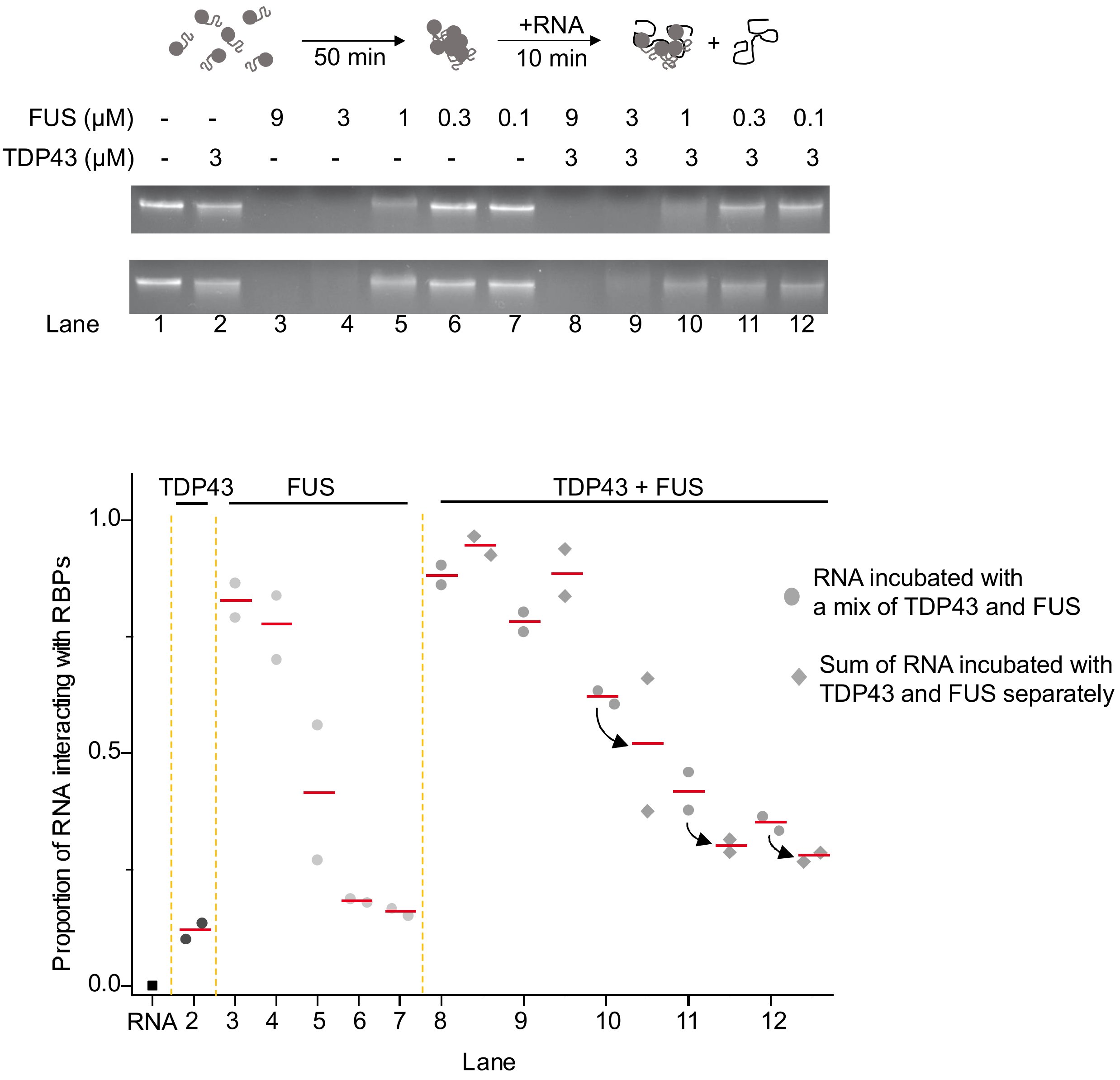


Supplementary Figure S8

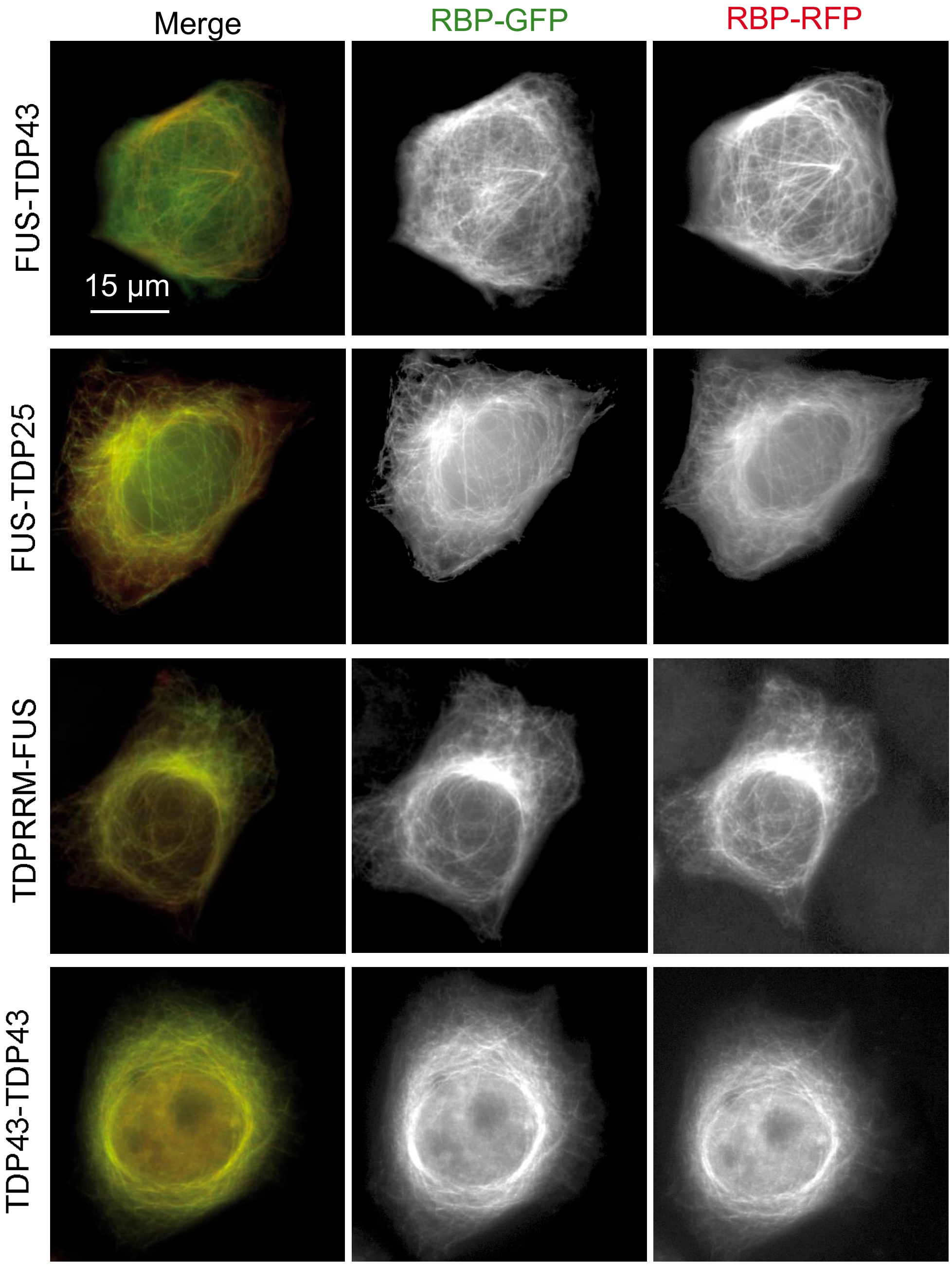


Supplementary figure S9

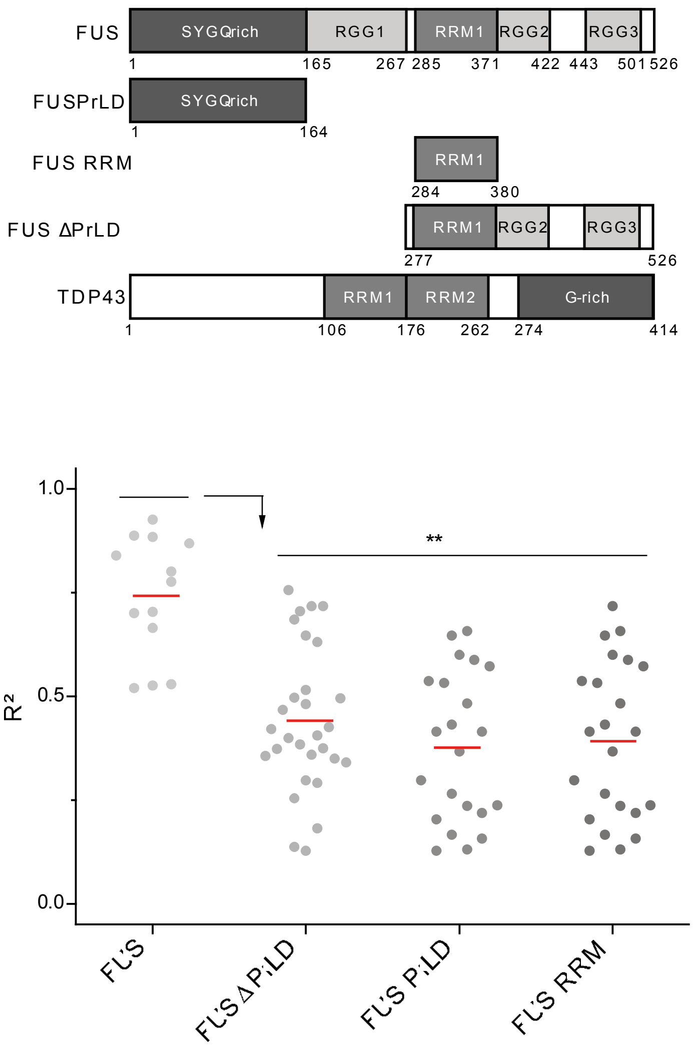


Supplementary figure S10

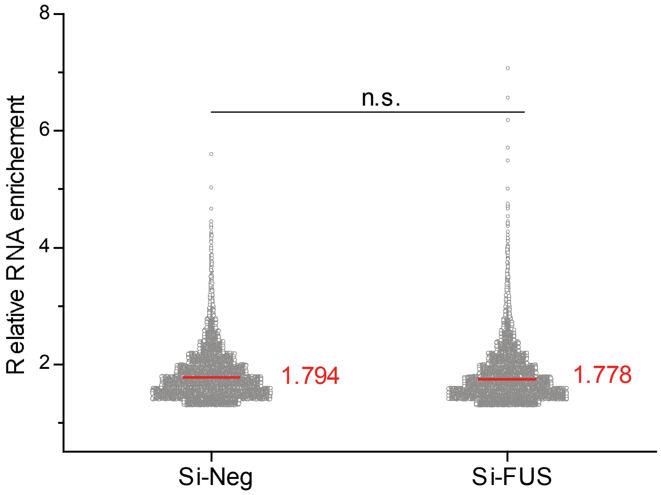


Supplementary Figure S11

Table of plasmids used for the Microtubule Bench bait-prey method

| Plasmid name | Sequence | tag | Primers or reference |
|---|---|---|---|
| FUS-RFP-MBD | FL (1-526) | RFP-MBD | [1] |
| TDP43-RFP-MBD | FL (1-414) | RFP-MBD | [1] |
| FUS-GFP | FL (1-526) | GFP | [1] |
| TDP43-GFP | FL (1-414) | GFP | [1] |
| YB1-GFP | FL (1-324) | GFP | [2] |
| Lin28-GFP | FL (1-209) | GFP | [2] |
| HuR-GFP | FL (1-326) | GFP | [1] |
| G3BP-GFP | FL (1-466) | GFP | [2] |
| Larp6-GFP | FL (1-491) | GFP | [2] |
| U2AF-GFP | FL (1-475) | GFP | [1] |
| Sam68-GFP | FL (1-443) | GFP | [3] |

Supplementary table 1

Table of plasmids used for the Microtubule Bench compartmentalization method

| Name | Sequence | Primers or reference |
|---|---|---|
| FUS-GFP-MBD | FL (1-526) | [1] |
| FUS-RFP-MBD | FL (1-526) | [1] |
| TDP43-GFP-MBD | FL (1-414) | [1] |
| TDP43-RFP-MBD | FL (1-414) | [1] |
| Sam68-RFP-MBD | FL (1-443) | [3] |
| HuR-RFP-MBD | FL (1-326) | [1] |
| TDP RRM-GFP-MBD | 98-263 | GCGTTAATTAAGCCACCATGGCAGTCCAGAAAACATCCGAT /GCGGGCGCGCCTGGTGCTTAGGTTCGGCATTGGA |
| TDP Nter-GFP-MBD | 1-102 | GCGTTAATTAAGCCACCATGTCTGAATATATTCGGGTAACCG/GCGGGCGCGCCTGTGTTTCTGGACTGCTCTTTTCAC |
| FUS RRM-GPF-MBD | 284-380 | GCGTTAATTAAATGAATTCAGACAACAACACCATC /GCGGGCGCGCCTTGCGAGTAGCAAATGAGACCTTG |
| FUS PrLD-GFP-MBD | 1-164 | GCGTTAATTAAGCCATGGCCTCAAACGATTATACCC/ GCGGGCGCGCCatACTGCTGCTGTTGTACTGGTT |
| FUS ΔLCD | 277-525 | [1] |
| TDP ΔPrLD-RFP-MBD | 1-277 | [1] |
| TDP25-RFP-MBD | 208-414 | GCGTTAATTAAATGCGGGAGTTCTTCTCTCAGTACGGGG/ GCGGGCGCGCCATCATTCCCCAGCCAGAAGACTTAGAA |
| TDP35-RFP-MBD | 102-414 | GCGTTAATTAAATGACGGAATATGAAACACAAGTGAAAG /GCGGGCGCGCCATCATTCCCCAGCCAGAAGACTTAGAA |
| TDP PrLD | 270-414 | [1] |
| mCherry-MBD | FL | [2] |

Supplementary table 2

Table of recombinant proteins (full length and truncated) produced for *in vitro* experiments.

| **Protein name** | **Tag** | **Sequence** | **Molecular Weight (kDa)** |
|---|---|---|---|
| **TDP43** | **6His** | **1-414** | **44** |
| TDP RRM | 6His-HA | 98-213 | 20 |
| TDP25 | 6His-HA | 208-414 | 25 |
| **FUS** | **6His** | **1-526** | **55** |

Table of plasmid for production of recombinant proteins (truncated forms).

| **Name** | **Restriction sites** | **Primers forward** | **Primers reverse** |
|---|---|---|---|
| TDP RRM | Nhe1/BamH1 | GCGGCTAGCGGCAGTCCAGAAAACATCCGA | GCGGGATCCGTGCTTAGGTTCGGCATTGG |
| | HindIII/Xho1 | GCGAAGCTTATGGGCTACCCCTACGACGTG | GCGCTCGAGGTGCTTAGGTTCGGCATTGG |
| TDP25 | Nhe1/BamH1 | GCGGCTAGCGCGGGAGTTCTTCTCTCAGTA | CGCGGATCCCATTCCCCAGCCAGAAGACTTAGA |
| | HindIII/Xho1 | GCG AAGCTTATGGGCTACCCCTACGACGTG | CTCGAGCATTCCCCAGCCAGAAGACTTAGA |

Supplementary Table 3

Table of primary antibodies used for immunofluorescence experiences

| **Antibodies** | **Species** | **Epitope** | **Reference** |
|---|---|---|---|
| α-TDP43 | Mouse | AA 1-260 (Nter) | mAB ABnova H00023435-M01 |
| α-TDP43 | Rabbit | (Cter) | pAB Protein Teck 12892 |
| α-FUS | Mouse | YGGQQQSYGQ (PrLD) | Novus CLO 190 mAB-03309 |
| α-FUS | Rabbit | AA 1-50 (PrLD) | Novus Bio pAB NB 100-565 |
| α-HA | Rabbit | Tag HA | H6908 Sigma |
| α-Myc | Mouse | Tag Myc | sAB1305535 Sigma (9E10) |
| α-Dig | Chicken | Tag Dig | pAb AB51946 Abcam |

Supplementary table 4

[1] Maucuer A, Desforges B, Joshi V, Boca M, Kretov D, Hamon L, et al. Microtubules as platforms for probing liquid-liquid phase separation in cells: application to RNA-binding proteins. Journal of cell science 2018.
[2] Boca M, Kretov DA, Desforges B, Mephon-Gaspard A, Curmi PA, Pastre D. Probing protein interactions in living mammalian cells on a microtubule bench. Scientific reports 2015;5:17304.
[3] Pankivskyi S, Pastre D, Steiner E, Joshi V, Rynditch A, Hamon L. ITSN1 regulates SAM68 solubility through SH3 domain interactions with SAM68 proline-rich motifs. Cellular and molecular life sciences : CMLS 2020.
